# ColTapp, an automated image analysis application for efficient microbial colony growth dynamics quantification

**DOI:** 10.1101/2020.05.27.119040

**Authors:** Julian Bär, Mathilde Boumasmoud, Roger Kouyos, Annelies S. Zinkernagel, Clément Vulin

**Affiliations:** Department of Infectious Diseases and Hospital Epidemiology, University Hospital Zurich, University of Zurich, Zurich, Switzerland; Institute of Medical Virology, University of Zurich, Switzerland; Institute of Biogeochemistry and Pollutant Dynamics, ETH Zurich, Zurich 8092, Switzerland; Department of Environmental Microbiology, Eawag, Dubendorf 8600, Switzerland

**Author notes:** These authors contributed equally to this work (Julian Bär, Mathilde Boumasmoud).

**Keywords:** application, microbial colonies, growth dynamics, image analysis

## Abstract

Phenotypic heterogeneity occurs in a population of genetically identical bacteria due to stochastic molecular fluctuations and environmental variations. In extreme cases of phenotypic heterogeneity, a fraction of the bacterial population enters dormancy, and these metabolically inactive or non-dividing bacteria persist through most antibiotic challenges. These subpopulations of persister cells are difficult to study in patient samples. However, the proportion of persisters in a sample can be accessed by physically separating bacteria on a plate measuring the time until colonies become visible as dormant bacteria resume growth later than their active counterparts and form smaller colonies.

Here, we present ColTapp (Colony Time-lapse app), an application dedicated to bacterial colony growth quantification, freely available for download together with its MATLAB source code or as a MacOS/Windows executable. ColTapp’s intuitive graphical user interface allows users without prior coding knowledge to analyze endpoint or time-lapse images of colonies on agar plates. Colonies are detected automatically, and their radius can be tracked over time. Downstream analyses to derive colony lag time and growth rate are implemented.

We demonstrate here the applicability of ColTapp on a dataset of *Staphyloccocus aureus* colony time-lapse images. Colonies on dense plates reached saturation early, biasing lag time estimation from endpoint images. This bias can be reduced by considering the area available to each colony on a plate.

By facilitating the analysis of colony growth dynamics in clinical settings, this application will enable a new type of diagnostics, oriented towards personalized antibiotic therapies.

## Introduction

Delayed and insufficient clearance of bacterial infections leading to treatment failure is associated with antibiotic resistance as well as antibiotic persistence. In antibiotic persistence, a subpopulation of bacteria, termed persister cells, can survive antibiotic challenges due to their phenotypic state of metabolic inactivity and then reconstitute the population by resuming growth when the stress is relieved (Balaban *et al*., 2004). This manifestation of phenotypic heterogeneity exists even in homogeneous liquid cultures, where each bacterial cell experiences the same local conditions (Ackermann, 2015). On top of this, bacteria often aggregate in dense communities such as biofilms, where environmental conditions are highly heterogeneous and further promote wide phenotypic distributions (Stewart & Franklin, 2008).

The clinical relevance of antibiotic persistence remains poorly understood. The main reason for this is that the existence and extent of persistence is difficult to assess because it is a transient phenotype and only concerns a small fraction of the bacterial population. However, these inactive bacteria are lagging longer than already actively dividing bacteria when plated on nutrient-rich agar medium, and the bacterial colonies’ macroscopic appearance time is a good proxy of single-cell lag time (Guillier *et al*., 2006, Levin-Reisman *et al*., 2010). For this reason, Levin-Reisman and co-authors proposed ScanLag, a platform to monitor colony growth dynamics with time-lapse imaging (Levin-Reisman *et al*., 2010, Levin-Reisman *et al*., 2014).

In clinical settings, the direct observation of colony growth dynamics from bacteria recovered from infection sites is rare (Barr *et al*., 2016, Vulin *et al*., 2018). Typically, in clinical microbiology laboratories the patient’s samples are plated and observed at a single timepoint, e.g. after 18 hours of incubation. Colonies which are smaller than the bulk at that timepoint have been associated with antibiotic tolerance, persister cells and relapsing infections (Proctor *et al*., 2006). Small colonies can result either from mutations that affect growth rate (small colony variants (Proctor *et al*., 1995, Kahl *et al*., 2016)) or from heterogeneous growth resumption (Jõers *et al*., 2010, Levin-Reisman *et al*., 2010, Vulin *et al*., 2018). Reading solely the colony size at a given timepoint does not allow to distinguish between these two phenomena. Moreover, while a bimodal classification, e.g. small or “normal”, may be appropriate to describe differences in colony size due to genetic changes (von Eiff *et al*., 2006, Guérillot *et al*., 2019), it fails to be quantitative when the range of observed colony sizes is not bimodal. This is a problem because phenotypic heterogeneity in single cell lag times typically results in unimodal colony size distributions with long tails of small colonies, and the experimenter is left with defining subjective cutoff values.

We anticipate that investigation of the antibiotic persistence phenomenon’s clinical relevance will greatly rely on colony growth dynamics analyses. Therefore, we propose to bridge the technical gap with a new analysis tool, the Colony Time-lapse application (ColTapp). Its goal is to promote wide access to colony growth dynamics quantification and enable diagnostic microbiology laboratories to further improve their routines. ColTapp is an image analysis pipeline embedded in an intuitive graphical user interface, which allows any user to derive colony sizes, growth rates and appearance times from time-lapse images of bacterial colonies. It also includes the possibility to estimate colony growth parameters from endpoint images. Additionally, it can report metrics characterizing color, shape and proximity of neighboring colonies. We speculate that ColTapp will prove useful in a broad spectrum of microbiology applications, such as species identification in environmental samples as shown by Ernebjerg & Kishony (2012).

A technical problem common to both clinical and environmental samples is the difficulty to assess their initial bacterial density: samples can be accidentally plated at high or uneven densities. Yet, colonies compete for nutrients on the agar plates and thus the total number of colonies and their spatial distribution impacts their size at an endpoint (Chacón *et al*., 2018). This results in difficult appearance time estimation. To address this problem, as well as demonstrate the applicability of ColTapp, we present a dataset of *Staphylococcus aureus* bacterial colony time-lapse images and explore their response to density. The colonies’ Voronoi cell areas, one of the spatial metrics computed by ColTapp and previously explored in the context of colony growth (Chacón *et al*., 2018), can be taken into account to minimize the bias in appearance time estimation due to density.

## Results

### A user-friendly application implementing image- and downstream analysis

ColTapp takes images as input and implements image analysis functions to detect microbial colonies and track colony radius over time when images are part of a time-lapse sequence. Common formats (png, tiff and jpeg) and either color (Red Green Blue) or grayscale images are supported. ColTapp also includes downstream analysis steps, to extract biologically relevant data such as colony growth rate and appearance time. A graphical user interface (GUI) provides access to all functionalities and displays the images, with the possibility to visualize certain data, such as the circles around the detected colonies (Fig. 1A, S1 and S2). The interface enables early visual evaluation of the results by including simple data visualization options (SI text 9). The generated data can be exported in a standard csv file (Fig. S3).

**Figure 1:**
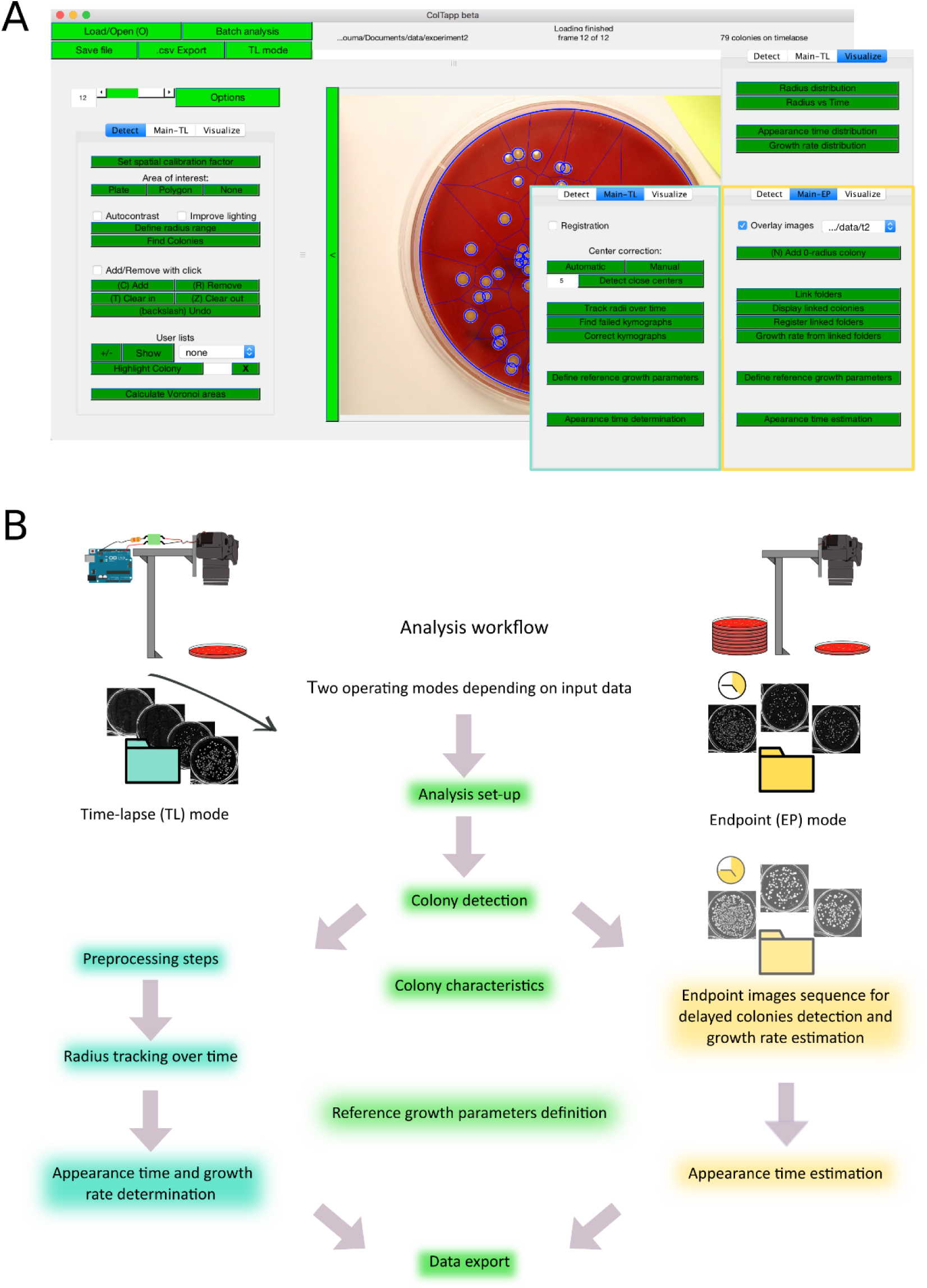
**A. Graphical user interface overview.** For visualization, results are overlaid onto the images: typically, the circles showing detected colonies and the edges showing the Voronoi tessellation shown here in blue on the image. The main analysis panel, on the left of the image, is separated in three tabs as depicted, allowing the user to detect colonies on a plate (*Detect* tab), analyze either time-lapse or endpoint images (*Main-TL* tab and *Main-EP* tab, contoured in turquoise and yellow respectively) and produce basic graphics from the obtained data. **B. Application workflow**. Schematic of a simple image acquisition setup including a camera holder. An Arduino board can be used to trigger the camera automatically for time-lapse imaging of a plate (Material and methods for implementation). ColTapp operates in two modes: *Time-lapse* (TL) and *Endpoint* (EP) mode depending on input data, illustrated by the two different folders (turquoise and yellow respectively). The turquoise highlighted functionalities are specific to the *Time-lapse* mode, while the yellow highlighted ones are specific to *Endpoint* mode. In the middle, the green highlighted functionalities are common to both modes.

The application’s workflow from raw images to exported data is illustrated in Fig. 1B. The program operates in two different modes, depending on the user input data: endpoint images or time-lapse images. In the first mode, *Time-Lapse* (TL), images in a folder are considered as an ordered time-lapse image series of a single plate. In the second mode, *Endpoint* (EP), all images in a folder are considered independent from each other. Typically, they are an aggregate of images from several plated samples or from replicates of the same sample, captured at a single timepoint.

In the following sections, implementation and performance of the two main image analysis algorithms (colony detection and radii tracking over time) are described, referring to the supplementary information for more detailed directives to the user concerning interaction with the interface, quality control functionalities and parameters optimization.

Then downstream analyses to derive growth parameters from the resulting data are demonstrated on a dataset of 22 time-lapse movies of *S. aureus* colonies plates, designed to include both homogeneous and heterogenous colony lag time and various colony densities. In addition to growth parameters, a palette of colony characteristics may be exported by the user for further analysis, including shape and color metrics and spatial metrics (describing the colony neighbors’ proximity, SI text 3).

In conclusion, an example is given to show how the ColTapp-derived measurements of radius, appearance time and spatial metrics can be used to assess the influence of colony density on appearance time estimation from endpoint images and how spatial metrics can be used to correct for density effects.

### Colony detection

#### Algorithm implementation

Colony detection uses a series of image analysis operations starting from a grayscale image (Fig. 2A) and depends on user-specified minimal and maximal expected radius of colonies (see SI text for additional pre-processing possibilities). ColTapp uses top-hat filtering with a disk-shaped structuring element based on the minimal expected radius to reduce lighting gradients and other inhomogeneities of the agar background (Fig. 2B). Local adaptive thresholding is then used to create a binary image (Fig. 2C). This image is cleaned from artifacts and unwanted objects by discarding small and big isolated objects (threshold defined by minimal and maximal expected radius, respectively). Objects with a low extent (foreground/background pixel ratio), are discarded as well (Fig. 2D). Next, distance transformation is used to derive local minima, which are subsequently filtered by minima imposition. Overlapping colonies are separated with watershed segmentation (Fig. 2E). Sequentially, ColTapp extracts images containing an isolated object (img_crop_) from the top-hat filtered grayscale image and further performs circular filtering and contrast enhancement (Fig. 2F). Objects are discarded if the proportion of low intensity pixels within img_crop_ is above a threshold defined by the median intensity of all objects. A two-step circular Hough transform is applied to img_crop_ by the *imfindcircles* function (Yuen et al 1990, Davies 2005) available in MATLAB to detect all probable colony circles within the given radius range (all circles in Fig. 2G). False positive circles may be detected (red circles in Fig. 2G) and sequential quality control functions are used to exclude these. Circles are excluded based on their distance from the image boundaries of img_crop_, proportion of pixels categorized as foreground within the circle, proportion of overlap with each other, distance between each other and relative fitting quality. After every img_crop_ is processed, a final quality control round is deployed to remove circles with extremely close centers (Fig. 2H). Several parameters involved in circle detection and quality control can be tuned to potentially improve performance (SI text 3.2). In addition, the user may intervene directly to correct the results (SI text 3.3).

**Figure 2:**
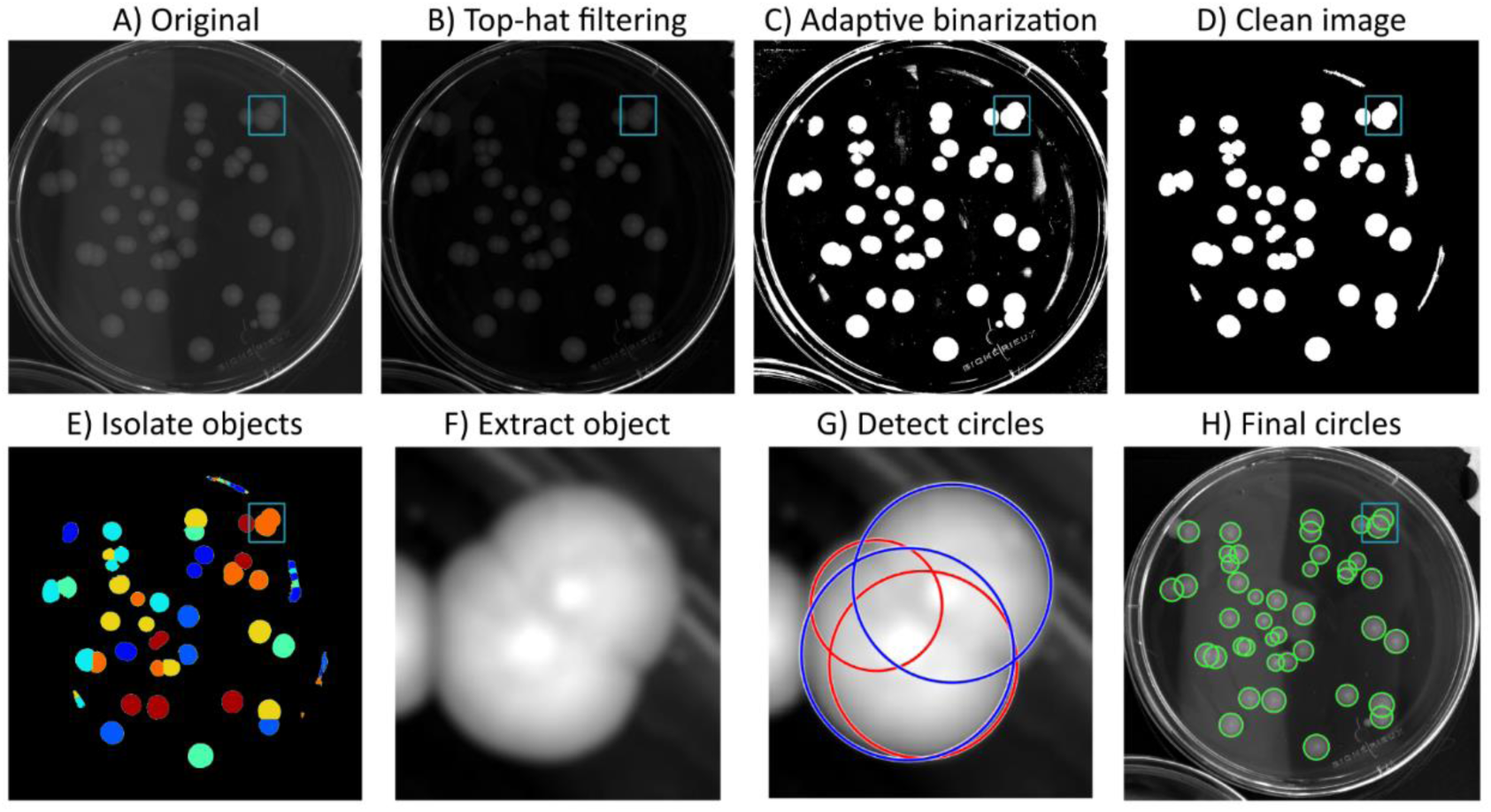
Colony detection. A series of image analysis steps are utilized to detect colonies on an image. **A**. Original grayscale image. **B**. Top-hat filtering of A. **C**. A binary image is obtained by adaptive local thresholding. **D**. Noise is filtered out of binary image. **E**. Distance transformation and watershed segmentation are used to identify isolated objects. **F**. Sequentially, each minimal sized image containing an isolated object is extracted. **G**. Potential circles are detected with a two-step Hough transform (all circles). A series of automatic quality control functions are used to discard false positive circles (red circles). Only circles passing the quality control are kept (blue circles). **H**. Final detected circles. The small rectangle in A to E and H marks the region of the extracted object shown in F and G.

In *Endpoint* mode, colony detection is done independently on each frame. In *Time-lapse* mode, it is done on a single frame, as the colony radius can subsequently be tracked over time (see below).

#### Algorithm performance

For the example shown in Fig. 2, a 1956×1936 sized grayscale image displaying 42 *Staphylococcus aureus* colonies after 70.5 h of growth on Columbia sheep blood with an average radius of 53 pixels, the computation from raw grayscale image to isolated objects took 1.57 s and the circle detection and quality controls took 0.98 s.

Computation efficiency and accuracy of the colony detection algorithm was tested with a selection of 26 differing images (Fig. S4) of the bacterial species *S. aureus, Staphylococcus epidermidis, Pseudomonas putida, Acinetobacter johnsonii* and *Alteromonas macleodii* as well as bacteriophage plaques on *Escherichia* coli (Pitol *et al*., 2017, Rodríguez-Verdugo *et al*., 2019) using the standard agar media necessary for optimal growth of the respective species (Colombia sheep blood, tryptic soy broth, lysogeny broth, marine broth, etc.). On this collection of images, which displayed colonies with an average radius of 28 pixels (SD = 15 pixels), colony detection took on average 0.049 s (SD = 0.033 s) per detected circle. Detailed values for each image are shown in Table S1.

Generally, ColTapp accurately detected colonies when the area of interest, expected radius range and the method for conversion of a color image into a grayscale image were set for each image separately (SI text 2.1). Note, that although colonies are usually lighter than the background, phage plaques are darker than the bacterial lawn: an option to find darker-than-background circles is available.

Overall, false positive rate varied substantially, ranging from 0% to 55.6% (median = 2.8%) (Fig. 3A). Almost all images with false positive rates exceeding 5% had most of their wrongly detected circles in clusters (marked with blue symbols in Fig. 3A), usually in areas with lighting artifacts. These clusters can be cleared with a few clicks and do not pose a problem in our opinion. The one other case with a high false positive rate was an image displaying colonies of the marine bacterial species *Alteromonas macleodii* and a lot of chitinous debris, forming particles which were only marginally different in size as compared to the bacterial colonies. Therefore, false positive rate should not exceed 5% if ideal imaging conditions are maintained. Additionally, we did not observe any correlation between false positive rate and total number of colonies on a plate (Fig. 3A).

**Figure 3:**
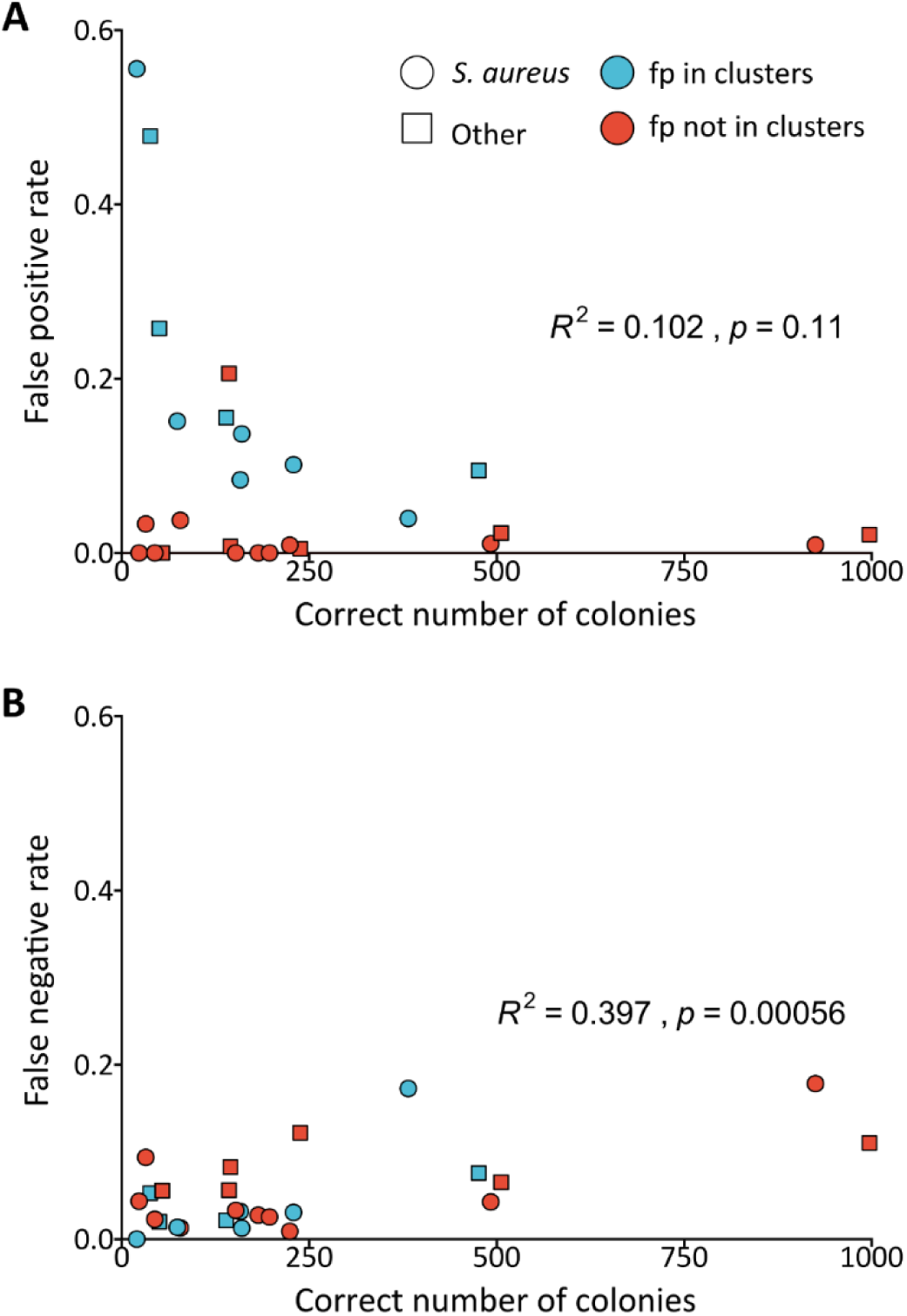
Accuracy of colony detection. A collection of 26 images of different species and agar media were used to benchmark the colony detection accuracy. **A**. False positive rate was not increasing with total number of colonies on a given image (Pearson correlation, p = 0.11). Some images had imperfect lighting conditions which resulted in glares and reflections. Most false positive circles were detected in clusters in these regions (blue circles, fp: false positive). **B**. False negative rate was correlated with total number of colonies (Pearson correlation, p = 0.00056). We did not observe an association between false negative rate and false positive clusters.

The false negative rate varied between 0% and 17.8% (median = 3.7%) and slightly increased with total number of colonies on the plate (Fig. 3B). Some of the images with high false negative rates had strong lighting gradients, high size heterogeneity or a high number of overlapping colonies all of which can decrease performance of ColTapp’s colony detection algorithm.

Note that computational speed is inversely proportional with image size, average colony size and number of colonies. A high proportion of plate area covered with colonies with high amount of overlap will generally yield poorer detection results and require manual correction (Table S1). In conclusion ColTapp successfully detects colonies from various bacterial species on distinctly colored agar plates. Adapting imaging conditions towards homogeneous light and reduced reflections can reduce the amount of manual correction required.

### Tracking colony radius over time

#### Algorithm implementation

In order to track colony radii over time, applying the previously described colony detection method on each frame would be slow and would require extensive manual correction since it is prone to misclassifications, especially at early timepoints when colonies are not yet visible. Therefore, ColTapp proposes to initially detect all colonies on a frame corresponding to a late timepoint, which is set as the *reference frame*. Once colonies have been detected on a late frame of the time-lapse series, their radius is tracked on the previous frames. For each time frame, ColTapp extracts sub-images of each colony from the main image. Intensity of each sub-image is scaled based on minimal and maximal intensity values of the last image of the time-lapse. These sub-images are then unwrapped with a polar transformation (Fig. 4A). The placement of the colony center position is important in this step since a misplaced center’s distance to the edge of the colony is not equal to the radius. Image drift correction as well as center correction functions are implemented in ColTapp to improve centering (SI text 5.2). Each transformed sub-image is averaged into a single intensity vector representing the intensity gradient from the colony center towards its edge. This averaging step reduces the noise introduced by slight off-centering of colonies and reduces file sizes. Finally, all vectors resulting from the corresponding sub-images of each time frame are combined into a kymograph, which represents the colony radial growth as a curve in intensity (Fig. 4A). The kymograph’s highest contrast edge corresponds to the colony radius over time. Since the presence of neighboring colonies would create blurry regions in the top of kymographs and thus hinder edge detection (Fig. 4B), ColTapp ignores the angle ranges corresponding to adjacent colonies for kymograph creation (SI text 6.1).

**Figure 4:**
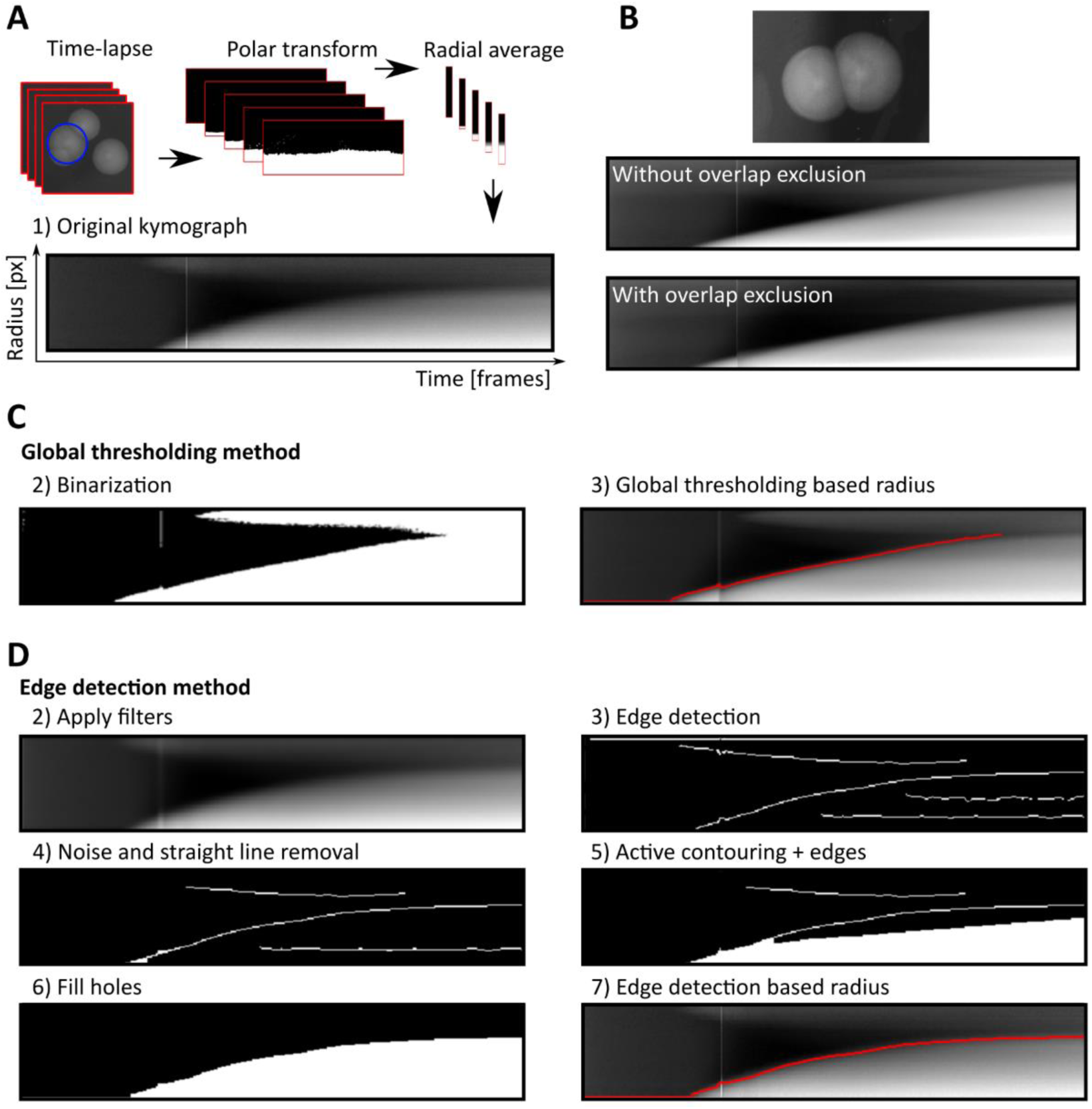
Time-lapse processing. **A**. A polar transformation is applied to grayscale sub-images of each colony and intensity values are averaged over the radius for each frame in the time-lapse. These radial average intensity vectors are combined into a kymograph representing colony radius growth. **B**. Overlapping colonies result in blurry regions in the top part of the kymograph. The angle range corresponding to overlapping colonies is excluded to reduce this effect. Kymographs can be processed either by **C**. *Global thresholding* or **D**. *Edge detection* method. **C**. In the *Global thresholding* method, a threshold is applied to a filtered version of the kymograph to derive a binary image. **D**. The *Edge detection* method uses a series of filtering steps, edge detection algorithms and morphological operations to optimize the binary image.

In order to detect the kymograph edge to derive colony radius over time, two methods are implemented: *Global thresholding* and *Edge detection*. Both methods first apply an automatic contrast enhancement function, then a pixel-wise adaptive low-pass Wiener filter (Lim, 1990) followed by a circular averaging filter (pillbox) to the kymograph to derive a smoother image. Additionally, a low-pass threshold for maximal intensity may be applied.

These pre-processed images are then further transformed with either a *Global thresholding* or *Edge detection* method to create binary images. The default *Global thresholding* method uses Otsu’s method (Otsu, 1979) to derive a global threshold from which a user-defined constant is subtracted to prevent biasing towards background (Fig. 4C).The *Edge detection* method uses a series of edge detection and morphological operations (Fig. 4D). In brief, the method by Canny (1986) is used for initial edge detection. Subsequently, small isolated foreground pixels are removed with area opening operations. Straight vertical and horizontal lines, typically originating from light artifacts and particles on the agar respectively, are removed. Two iterative morphological closing operations are used with line- and disk-shaped kernels, respectively. Additionally, active contouring using the Chan-Vese method (Chan & Vese, 2001) is applied to the pre-processed kymograph to derive a second binary image with a filled region of probable colony area. The two binary images are merged, and morphological closing is performed to close potential gaps to successfully fill connected regions afterwards. A final morphological opening is applied to remove jagged edges. The fully connected object located in the lower right side of the kymograph (corresponds to end of time-lapse and close to colony center) is kept as the only foreground object in the binary image.

ColTapp includes automatic error detection and manual correction tools for the kymograph derived radius growth curves (SI text 6.2, Fig. S5).

#### Algorithm performance

Computational time was assessed on a subset of 10 time-lapse sequences (each of 410 or 423 frames) from our demonstration dataset. Total required time was proportional to the number of frames and colonies to process, because each frame and colonies are processed sequentially (Table S2). Therefore, computational time per colony and per frame is most representative of the computational efficiency of the colony radius tracking algorithm (average = 0.079 s, SD = 0.047 s). Additionally, the computational time was observed to increase with the size of the colonies on the *reference frame*, as the sub-images used for radius tracking are bigger.

We assessed the accuracy of the algorithm by manually evaluating the quality of the 1411 kymograph derived radial growth curves. The *Global thresholding* method usually yields correct binary images except for complex kymographs resulting from high amount of colony overlap and/or lighting artifacts. When using the default *Global thresholding* method to derive radial growth curves, some curves of our dataset were classified as incorrect (mean = 21%, SD = 15%) (Table S2). ColTapp inbuilt automatic quality assessment function detected on average 83% (SD = 9%) of these incorrect growth curves (Table S2). The *Edge detection* method is more complex and is suggested when the default *Global thresholding* method fails. Switching to the *Edge detection* method for the incorrect growth curves resulted in a reduction of the number of growth curves requiring further manual correction to 3.1% (SD = 2.7%) (Table S2).

### Appearance time and growth rate determination from time-lapse images

Tracking the colony radius over time using time-lapse imaging makes it possible to directly determine colony appearance time and growth rate. Colonies formed by non-swimming bacteria pass through three growth phases (Fig. 5A). Initially, when all bacteria forming the colony can access nutrients and thus contribute to its expansion, the colony radius increases exponentially while the colony also increases in height. Eventually, bacteria at the center of the colony do not have access to nutrients diffusing from the edge of the colony. The zone of growth at the edge of the colony becomes constant and the radial growth rate becomes linear (at this point radius = R_lin_, see Fig. 5B) (Pirt, 1967). Finally, when the nutrients become scarce, or when growth byproducts reach an inhibitory concentration, the colony enters the saturation phase, during which the radial growth rate starts decreasing until plateauing (Be’er *et al*., 2009). Fig. S6 shows the exponential phase of *S. aureus* colonies grown on sheep blood agar captured by time-lapse imaging with a macro lense. However, in typical settings where the entire plate is captured, the resolution is not high enough to observe the exponential phase. We define a detectable size threshold (R_thresh_), and the time at which a colony reaches this threshold as the *appearance time* (t_app_). The minimal possible R_thresh_ depends on the image quality and needs to be set at the same value for comparisons of t_app_ in different experiments. In our analysis setting, we assume that colonies reaching this size are already in the linear growth phase (R_lin_ < R_thresh_, Fig. 5B, C).

**Figure 5:**
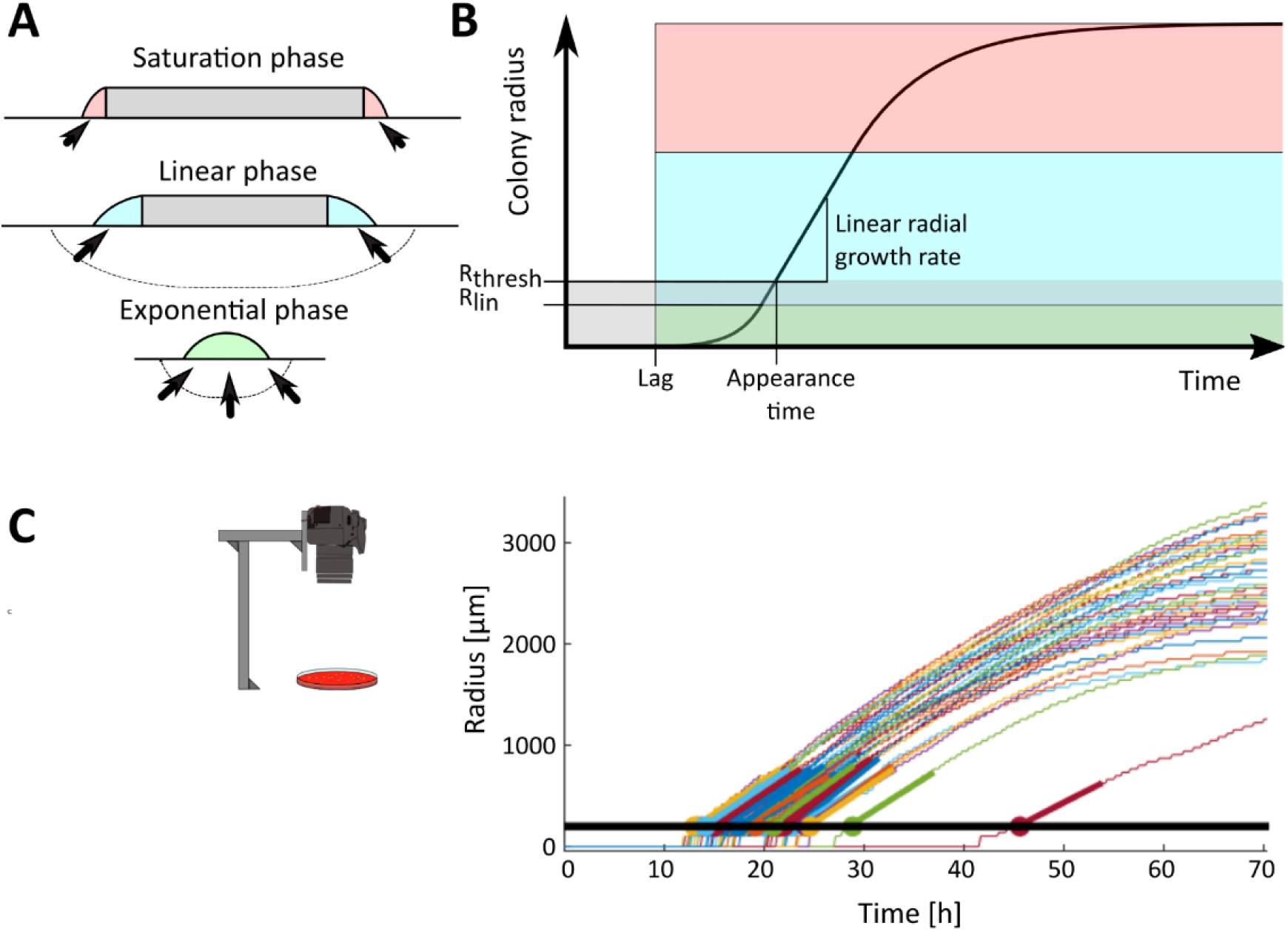
Colony radial growth curves. **A**. Schematic cross section of a colony in the three growth phases. The 3 colors represent the phases (green: exponential phase, blue: linear phase, light red: saturation phase). The grey depicted areas represent bacterial cells which do not contribute to colony expansion. The arrows represent the flow of the diffusing growth substrate, steady during the linear phase and decreasing during the saturation phase. **B**. Typical radial growth curve, with the three phases highlighted with colors corresponding to A. R_lin_ is the radius at which the radial growth rate becomes linear. The grayed zone indicates the size range at which a colony is not macroscopically visible (radius < R_thresh_). **C**. The typical radial growth curves obtained by time-lapse image analysis in settings where the whole plate is captured (represented by the camera holder and agar plate icon). On the graph, the black horizontal line represents the threshold radius (r_thresh_) used for appearance time calculation and thick colored line segments represent timespan used for linear regression to estimate appearance time. Appearance time is marked with bold circles.

ColTapp determines appearance time and growth rate by detecting the first of 6 consecutive frames (to avoid noise) for which a colony radius is bigger than R_thresh_ (default: 200 µm). From the detected frame, the radius values measured on the following frames, corresponding to a user-specified timespan in hours (Fr_lin_, default: 10 h), are used for a linear regression to determine linear radial growth rate and the exact time a colony reached R_thresh_ (Fig 5C). Fr_lin_ might need to be reduced in order to only use a timespan in which radial growth is approximately linear. Additionally, R_thresh_ can be adjusted depending on the image resolution and bacterial species investigated. The reason to perform this linear regression to determine appearance time instead of simply observing the time when a colony radius reaches R_thresh_ is that it reduces artificial noise, to be expected if camera resolutions are low (for example if 1 pixel = 50 µm).

Growth parameters such as the duration of the exponential phase and the colony linear radial growth rate depend on strain characteristic properties such as replication frequency, metabolic yield (Pirt, 1967), cell-to-cell adhesion (Nguyen *et al*., 2004) and surfactant production (Angelini *et al*., 2009). In addition, growth dynamics depend on the nutrient medium and physical properties of the agar surface such as substrate roughness (Gralka & Hallatschek, 2019) or water activity (Stecchini *et al*., 2001).

Therefore, for downstream analyses, reference growth parameters need to be measured with a control dataset (SI text 7). The reference lag time is the minimal time to colony appearance and can be observed by plating exponential phase bacteria which show little to no delay in growth resumption (Vulin et al. 2018). The reference linear radial growth rate is the maximal growth rate (GR_max_) observed in any experiment before competition between neighboring colonies kicks in and affects it.

The average colony growth rate derived from our control dataset (plates with less than 150 colonies) is 60.4 µm/h (SD = 4.7 µm/h). We define GR_max_ taking the 95^th^ percentile (65.0 µm/h) of the growth rate distribution.

### Appearance time and growth rate estimation from endpoint images

In some laboratories, setting-up high-throughput time-lapse imaging might not be possible, thus ColTapp proposes to estimate colony lag time from endpoint images. This is possible because when bacterial cells with a lag resume growth, they rapidly reach their maximal replication rate and the resulting colonies have growth rates similar to those of colonies emerging from cells with no lag (Vulin *et al*., 2018). One can thus estimate the colony appearance time based on colony radius R at a given timepoint (t_f_) by simple linear regression:

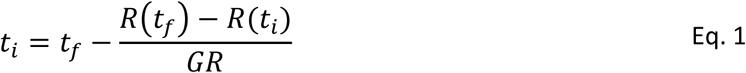

with R_thresh_ defining the radius (R) at time t_i_ = t_app_ and a reference growth rate (GR_max_).

Note that using a linear regression with a reference growth rate to estimate the appearance time has two strong assumptions: first, that all colonies are still in the linear phase of their growth and second, that they all have the same linear radial growth rate.

To allow users to evaluate the validity of these two assumptions, ColTapp proposes a functionality to analyze a sequence of endpoint images (SI text 8.2). By taking endpoint images at multiple timepoints and performing colony detection at each timepoint one can create a timeseries of colony radius. We exemplify this functionality by analyzing two series of 4 frames extracted from time-lapse image sequences of the demonstration dataset (Fig. 6A). ColTapp computes the slope between colonies’ radii at different timepoints (Eq. 1), which results in an individual colony radial growth rate for each time interval (Fig. 6B).

**Figure 6:**
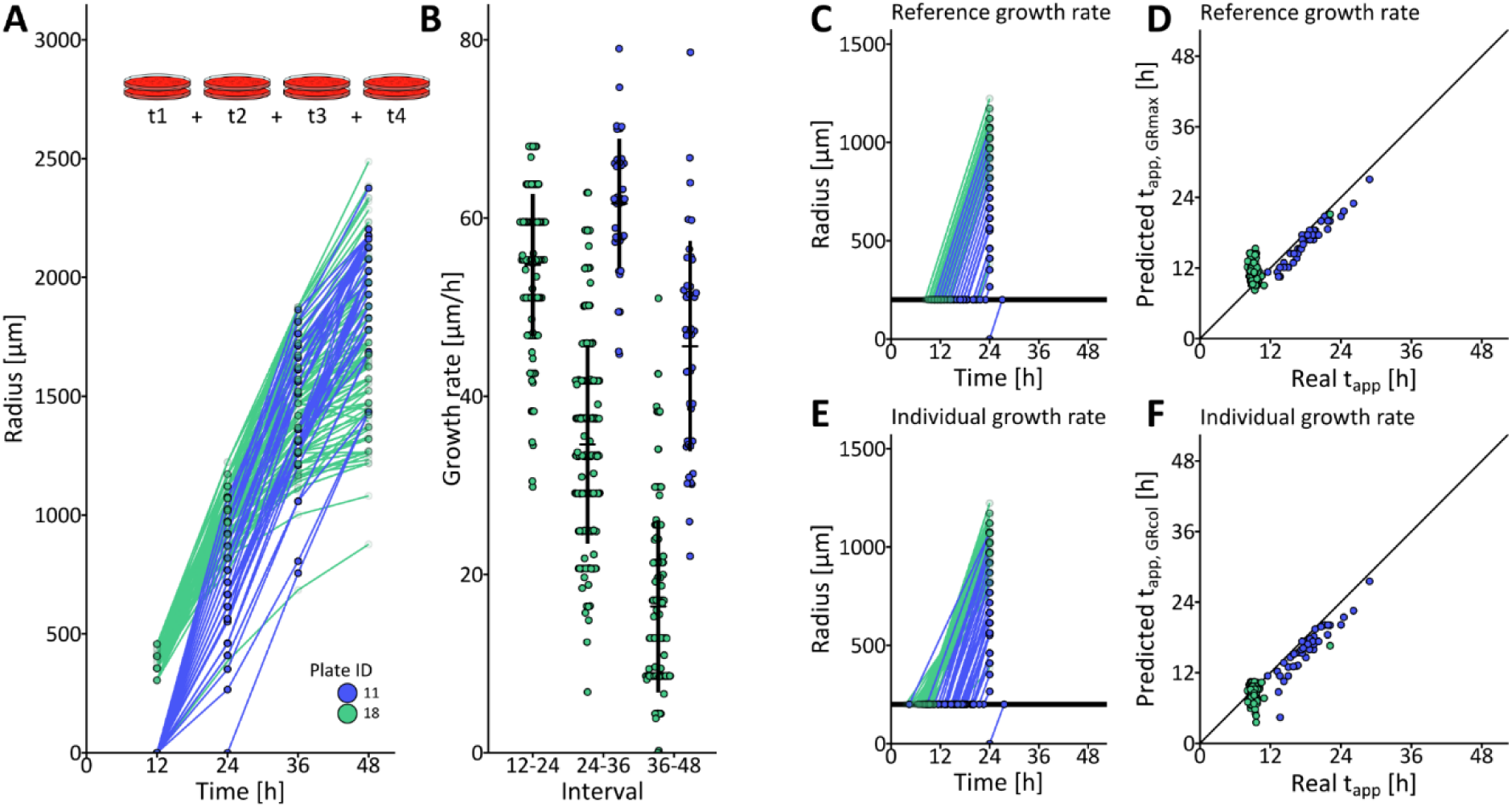
Growth rate and appearance time estimation from endpoint images. **A**. Four images corresponding to the timepoints 12 h, 24 h, 36 h and 48 h were extracted from the full time-lapse image sequences of two independent plates and separately analyzed. Color indicates plate ID. Each dot represents radius of one colony at the given timepoint and all dots corresponding to the same colony are connected by lines. **B**. Growth rate was estimated for each of the three intervals between the four timepoints for each colony. Crossbars indicate mean ± SD per timepoint and plate ID. When the radius is null at one of the timepoints involved in the interval, growth rate is not calculated. Therefore, no growth rate values are shown for Plate ID 11 at the first interval. **C. E**. Estimation of appearance time from radius at the 24h timepoint by linear regression. **D. F**. Comparison of the appearance time (t_app_) determined by the full time-lapse analysis (Real t_app_) and the appearance time estimated by linear regression (Predicted t_app_) **C. D** with a reference growth rate GR_max_ of 65 µm/h. **E. F** with individual colony growth rate.

This enables assessment of the entry time to saturation phase, identified by a decrease in growth rate. Moreover, should colonies have different growth rates, the individual colony radial growth rate can be used to estimate the appearance time, rather than a fixed reference growth rate for all colonies (Fig. 6C, D, E and F). Although using individual colony growth rates increased overall agreement between appearance times determined from time-lapse images and appearance times estimated from endpoint images (SI Fig. 7), it could not accurately predict appearance time for colonies past the linear phase (i.e. colonies on plate 18 at 24h in Fig. 6).

**Figure 7:**
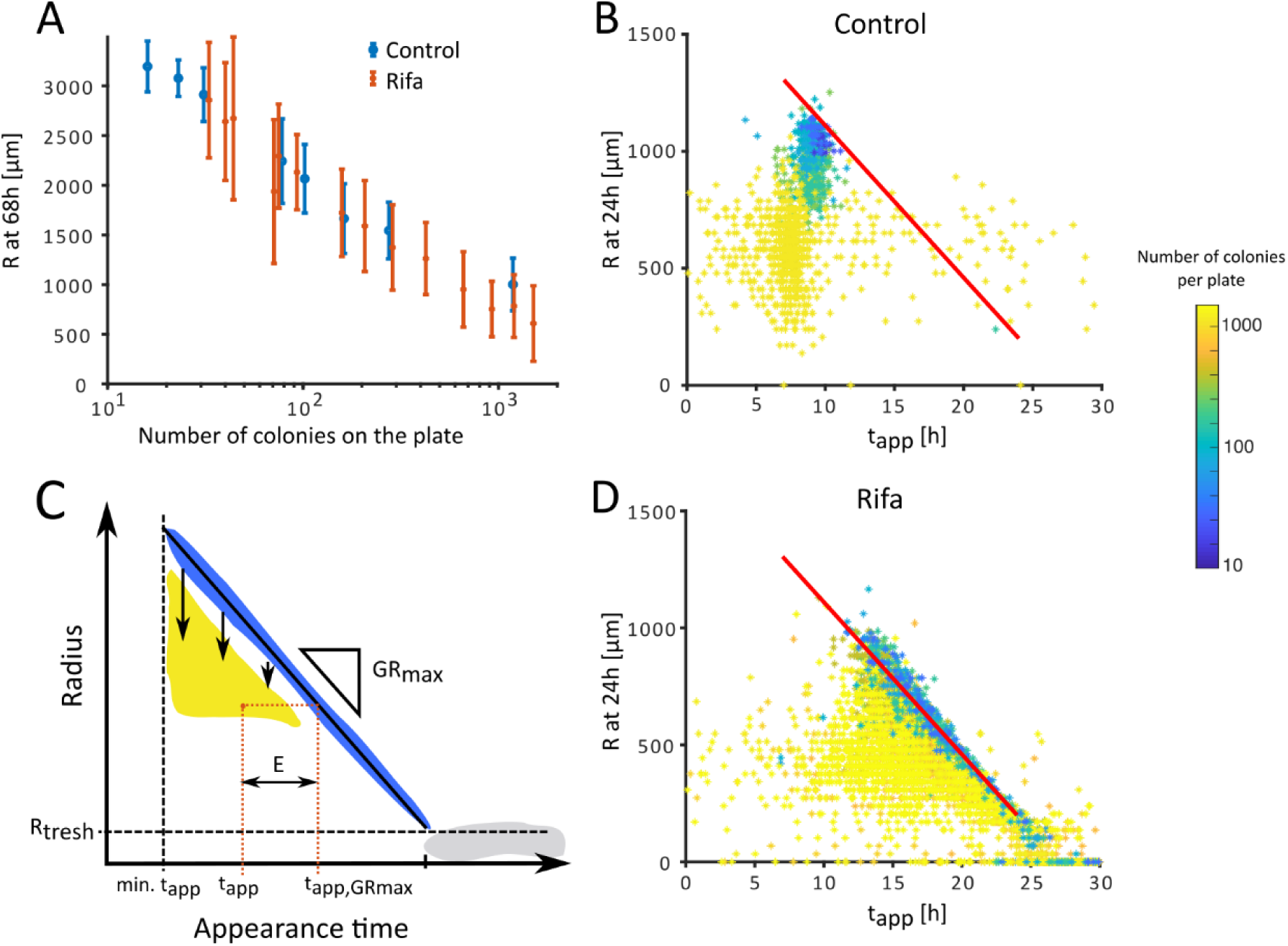
Effect of density. **A**. Since colonies compete for the limited resources of the agar plate, at a given time (here 68h), colonies on a dense plate have a smaller radius than colonies plated at low density. **B**. In our control dataset, the linear relationship expected between the colonies’ appearance time and their radius (red line defined by GR_max_ = 65 µm/h) was lost when colonies grew on dense plates (here with warmer colors) and was heterogeneous within a plate at a given time. Here, at 24 h, while some colonies were still following a linear growth, other colonies from the same plate had already entered saturation phase. **C**. This relationship between radius and appearance time at a given time can be visualized as a drop of the colony radius (yellow) from the ideal growth line (blue) as soon as a colony enters the saturation phase. An example of one colony is shown, represented by the orange dot. Its appearance time (t_app_) is given by its x-coordinate, and the appearance time estimated using a linear regression with GR_max_ is given by the x coordinate of its projection on the ideal growth line (t_app,GRmax_). The error (E) is the difference between these two values along the x-axis. Colonies that are below R_thresh_, and therefore did not appear yet fall in the gray area. **D**. The effect of plate density was also visible in our dataset of *S. aureus* exposed to rifampicin (abbreviated rifa on figure and further in text), with a larger variance in colonies’ appearance times and radii.

Note that using individual colony growth rates for appearance time estimation is appropriate to correct for differences in size independent from saturation, for example when a subpopulation on the plate has mutations affecting growth rate or mixed-species plates. Should colonies be past the linear growth phase, one needs to take saturation into account. We introduce an approach to density-based corrections of saturation in the next section.

### Response to density

One typical problem for environmental or clinical samples is that the number of colony-forming-units per milliliter of sample is often unknown before plating, resulting in unpredictable colony densities on the plates. Yet, colony growth dynamics are affected by the presence of neighboring colonies, which compete locally for nutrients and are affected by the total number and proximity of colonies on the plate (Dervaux *et al*., 2014, Chacón *et al*., 2018). As soon as the effects of competition with neighboring colonies become strong enough, a colony enters the saturation phase of its growth, meaning that its radial growth rate decreases over time until plateauing (Fig. 5). Therefore, colonies on denser plates are expected to be smaller than colonies on less dense plates at a given point (Fig. 7A).

On top of this, if the plating is heterogeneous, not all colonies have access to the same amount of resources. They therefore differ in size within a plate as soon as they enter the saturation phase. Thus, experiments aiming to compare either colony size or appearance time are biased by colony density.

We explored the response to density using our purposely designed demonstration dataset of colonies formed by the *S. aureus* lab strain Cowan I growing on Columbia sheep blood agar. Of the 22 time-lapse movies, eight were plated from a liquid culture at exponential phase to obtain a homogeneously growing population (control dataset, Fig. 7A). The other fourteen were plated after exposure to the antibiotic rifampicin, which arrests bacterial cell growth by stopping protein synthesis, resulting in a bacterial population with a wide appearance time distribution (further referred to as rifa dataset, Fig. 7A) (Kwan *et al*., 2013). In order to observe the impact of density on colony radius over time and elaborate a method to correct this bias, the replicates were plated at different densities over two orders of magnitude (ranging from 16 to 1509 colonies per plate) (SI Fig. 8, 9).

**Figure 8:**
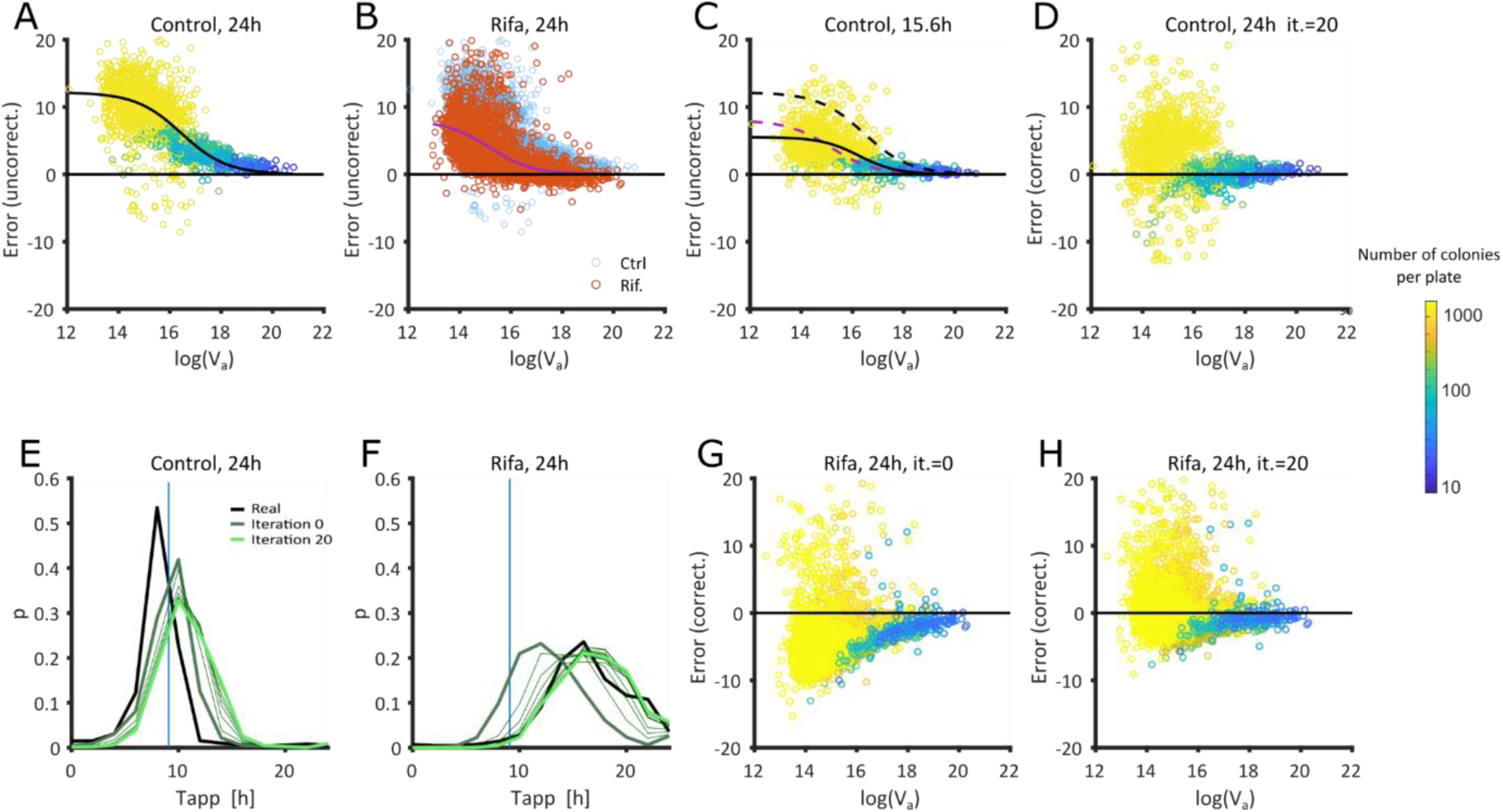
**A**. For a given timepoint (here 24 h, control dataset), the error between appearance time estimated by linear regression and appearance time determined by the full time-lapse analysis correlates with the Voronoi cell area (V_a_) and can be fitted using a logistic model (black line). Each dot represents one colony, colored by density (log values on the color bar, left side of figure) same as in C, D, G, H. **B**. The relation E(V_a_) observed at 24 h on the control dataset (blue dots underlaid on this plot) depends on the median lag time. Here it can be seen that E(V_a_) observed at 24 h on the rifa dataset (red dots, fitted by the purple line) is systematically smaller. **C**. Here the relation E(V_a_) observed at 15.6h on the control dataset is shown. The fit of this data (black solid line) is close to the fit of E(V_a_) observed at 24 h on the rifa dataset (dashed purple line). Dashed black line it the fit of E(V_a_) observed at 24 h on the control dataset. **D**. The result of the iterative correction for the control dataset observed at 24 h (shown in A before iterative correction). **E - F**. Visualization of the improvement of appearance time estimation by iterative corrections depending on time difference between median estimated t_app_ and the known median t_app_ of the control dataset (vertical blue line), yielding predictions closer to the appearance time determined by the full time-lapse analysis (real, bold black line). The initial guess (Iteration 0, bold dark green line) is corrected (thin lines) over 20 iterations to yield final corrected t_app_ (Iteration 20, bold light green line). Histograms are binned in 2h bins. **G-H** Estimated appearance time of the rifa dataset (shown without correction in B) is either corrected using E(V_a_) observed at 24 h on the control dataset (no iterations), or through 20 iterations (abbreviated it. on figure).

The effect of high densities on the colony growth rate is shown on Fig. 7BD displaying the colonies’ radii against their observed appearance times at a given timepoint. One may follow the evolution of this relationship over time (SI movie 2). At low densities, colonies grow at the maximal linear radial growth rate, GR_max_, which depends on strain and culture conditions (Fig. 5). Colonies that deviate from the GR_max_ appear below the expected linear regression line (Fig. 7C). Therefore, when performing a linear regression using GR_max_, the appearance time of colonies which are already in the saturation phase are overestimated (Fig. 6D). The systematic error (E) introduced by saturation corresponds to the difference between t_app, GRmax_ (the result of the linear regression with GR_max_) and the appearance time (t_app_) determined by the full time-lapse analysis (Fig. 7C):

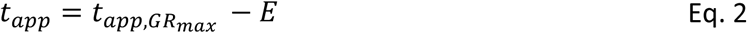

### Density correction

Chacón *et al*. (2018) proposed that the amount of nutrients available to each colony on the agar plate can be approximated with Voronoi cell areas, obtained by drawing equidistant lines between neighbor colonies centers. The correlation between the radius at the time of observation and colony Voronoi cell area is time dependent and strongest at a late growth stage, as colonies approach their maximal size (Chacón *et al*., 2018).

Aiming to use the Voronoi cell area (V_a_) to approximate the error in appearance time estimation introduced by saturation (E), we evaluated the mathematical relation between log(V_a_) and E in the control dataset. At 24 h, E was close to 0 for colonies with large V_a_ and increased for colonies with smaller V_a_ (Fig. 8A). The relation E(V_a_) was fitted with a modified logistic model (Fig. 8A):

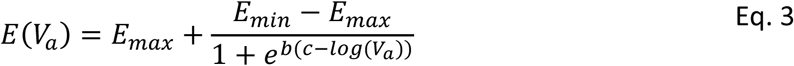

where E_min_ is set to 0, and E_max_, b and c are fitted parameters using the least square method. E_max,_ exists because even for infinitely small values of V_a_, colonies still grow by using nutrients immediately below them. b and c are shape parameters for the transition from E_min_ to E_max_.

As expected, the relation E(V_a_) was time dependent. When fitted through time (SI movie 3, SI Fig. 10A, B, C), a linear increase of E_max_ was observed starting as soon as the first colonies enter saturation phase. In addition to being time-dependent, E(V_a_) was observed to depend on the general lag time of the plate: at a given timepoint, colonies originating from populations with a large median lag time (e.g. rifa dataset) were in an earlier phase of their growth as compared to colonies originating from populations without lag (e.g. control dataset) (Fig. 8B). Ideally, one would adapt the relation E(V_a_) to take into account the median lag time of the plate, so that the correction applied to the rifa dataset observed at a given timepoint (t_obs_) corresponds to E(V_a_) fitted on the control dataset at an earlier timepoint, when the colonies were still at that stage of growth.

The difference in appearance time between the control dataset (median appearance time: 9 h) and the rifa dataset (median appearance time: 17.4 h) was Δt_app_ = 8.4 h. The correction applied to the rifa dataset when observing colonies at t_obs_ = 24 h should thus correspond to E(V_a_) fitted on the control dataset at t_obs_-Δt_app_ = 15.6 h (Fig. 8C). However, the median appearance time is not known *a priori* when observing colonies on a plate at an endpoint.

Therefore, we used an iterative process to estimate appearance time. An initial guess of the appearance time (t_app,i_) was obtained with Eq. 2 where E(V_a_) was fitted on the control dataset at t_obs_. Then the difference between the known median t_app_ of the control dataset and the median t_app,i_ of the observed plate (Δt_app,i_) was calculated. A new t_app,i_ was estimated with Eq. 2 where E(V_a_) was fitted on the control dataset at t_obs_ - Δt_app,i_, which in turn gave a new Δt_app,i_. This process was iterated until stabilization for a final estimate of t_app_ (Fig. 8E to H, Fig. S10 A to H and L).

Note that the shown approach tended to overcorrect the appearance time of colonies on dense plates due to the nature of the data: fully saturated colonies have little information left regarding their appearance time since the radius does not change with time anymore. Thus, this correction cannot perform well if colonies have reached late saturation phase (SI Fig. 10E to F). For this reason, in populations with large variance of appearance time, one single image may not allow a proper appearance time estimation, especially if some colonies stop growing before the late ones appear. In such cases, we recommend using several pictures of the same plate at different timepoints in order to minimize possible biases in estimations of lag time distributions.

## Discussion

We developed ColTapp, a user-friendly application with a graphical interface. This application extracts biologically relevant data from images of microbial colonies including size, growth rate and appearance time as well as color, shape and spatial metrics. Colony growth dynamics parameters are derived from time-lapse images. As it might not always be feasible to utilize high-throughput time-lapse imaging, ColTapp also includes a framework to estimate appearance time from endpoint images. Colony size at an endpoint is affected by colony density, resulting in a bias in appearance time estimation. Thus, we propose an iterative correction utilizing local plate density to moderate this bias. The correction is based on a control dataset and applied to a dataset with different growth characteristics (generalized long lag time).

In the past decades, the existence of phenotypic heterogeneity in bacterial populations has gained interest. Mathematical modeling of bacterial population dynamics in their ecosystems used to rely on average behavior. Advances in single cell microscopy and the development of models that can take into account the phenotypic state of thousands of individual bacteria shed light on the ecological function of phenotypic heterogeneity (Lennon & Jones, 2011, Ackermann, 2015). In medicine, while most bacterial infections can successfully be treated with antibiotics, treatment failure occurs even if the bacteria are not resistant to the antibiotics that are given. The existence of subpopulations unresponsive to treatment is thought to ultimately be the cause for these treatment failures.

Under standard laboratory conditions, the established methods to investigate phenotypic heterogeneity in bacterial populations are single cell microscopy and flow cytometry, but these methods are often expensive and technically difficult to setup and maintain. Moreover, many medical and environmental samples are too complex to study with these methods, either because they consist of complex substrates (e.g. viscous fluids) or they contain contaminating particles that may hinder proper quantification. The high fluctuation in quality and bacterial load in natural samples might also not justify the time and money investment required to carry those experiments out. In the case of microscopy, rare phenotypes are difficult to quantify if the setup does not allow the observation of large number of bacteria. Therefore, plating bacteria on nutrient media offers a cheap and easy way to assess lag time heterogeneity, as previously proposed (Guillier *et al*., 2006, Levin-Reisman *et al*., 2010). While time to growth resumption on undefined nutrient media reflects the dormancy state of the bacterial cells upon plating, one may be interested in following time to growth resumption of an active population upon plating on selective media. Indeed, this can be an opportunity to investigate the ability of a population to switch metabolic states (e.g. to a new carbon source) (Solopova *et al*., 2014).

Note that although colony formation can be seen as a good alternative to single cell behavior assessment, it is not a direct single cell observation. For example, clumped bacteria or an inhomogeneous spreading of cells on a surface, resulting in deposition of aggregates of cells instead of single cells on the agar, will likely result in an underestimation of single cell lag time. This needs to be kept in mind even when calibrating data to optimize analysis. Assessing the effect of the studied condition on clumping of bacteria might help avoiding false discoveries.

With ColTapp, we propose an easy to use graphical user interface, so that any laboratory, regardless of the technical equipment and image analysis expertise accessible, can approach single cell growth phenotypes by monitoring colony growth dynamics. We qualitatively compare ColTapp with other tools available in Table 1 below. Usage of colony size determined by image analysis to assess stressor effects is a long-established method (Dykes, 1999) and various end-user applications to facilitate colony counting and size assessment were developed over the years (e.g. a standalone GUI by Clarke *et al*. (2010) or an ImageJ plugin by Cai *et al*. (2011)) of which OpenCFU is one of the best known comprehensive colony counting and size assessment applications (Geissmann, 2013). Image analysis algorithms implemented by the various applications are usually similar and consist of a combination of pre-processing steps to clean raw images, a variety of thresholding algorithms to derive binary images and optional watershed segmentation to finally count and measure isolated objects. Variations exist with for example multiple rounds of pre-processing (Brugger *et al*., 2012) or a two-step colony detection algorithm for plate border and center (Chiang *et al*., 2015). Modern methods are using smartphones to acquire and process images all in one simple app (Wong *et al*., 2016, Austerjost *et al*., 2017) or are starting to use the power of deep learning algorithms (Andreini *et al*., 2018). All these applications are meant to analyze endpoint images, which do not capture colony growth dynamics. Levin-Reisman *et al*. (2010) released the ScanLag code package to analyze colony time-lapse images. Yet, although multiple groups use this type of analysis for their research (Ernebjerg & Kishony, 2012, Barr *et al*., 2016, Chacón *et al*., 2018, Liu *et al*., 2020), there is no comprehensive and flexible application including a graphical user interface currently available for this purpose. Therefore, by bringing together the standard endpoint analysis with time-lapse image analysis, our user-friendly ColTapp fills a gap in the collection of existing dedicated image analysis tools.

**Table 1:** Selected publications presenting colony images analysis tools. This selection of publications aims to display at one glance previous reports presenting tools and applications dedicated to image analysis of pictures of bacterial colonies grown on solid agar. We summarize the availability of code and GUI, and which data regarding colony characteristics are provided as well as what feature of each publications made them unique at the given publication time. For details and more publications, see main text.

|  | Availability |  | Data extracted from colonies on images |  |  |  |  | Unique feature |
| --- | --- | --- | --- | --- | --- | --- | --- | --- |
|  | Code | GUI | Count | Position | Size | Growth kinetics | Additional characteristics |  |
| Levin-Reisman et al. 2010 (ScanLag) | Yes | No | Yes | Yes | Yes | Yes | None | Analysis of colony time-lapse images |
| Geissmann 2013 (OpenCFU) | Yes | Yes | Yes | Yes | Yes | No | Shape and color metrics, number of adjacent colonies | Colony counter application |
| Wong et al. 2016 (APD Colony Counter App) | No | Yes | Yes | No | No | No | None | Smartphone colony counter application |
| ColTapp | Yes | Yes | Yes | Yes | Yes | Yes | Shape and color metrics, spatial metrics | Application for the analysis of colony time-lapse and endpoint images |

As for any image analysis tool, the quality of the analysis will highly depend on the quality of the images themselves. We recommend users to maintain homogenous lighting, avoid light flares and reflections and create good contrast using background modifications and proper focusing. ColTapp is flexible enough to allow processing of colored and grayscale images acquired either through dedicated platforms or with simple camera solutions. For endpoint images, this can be best achieved using a dedicated white box with diffused, indirect light, but recent phone cameras can also give decent results on a bench.

We propose an analysis of the linear radial growth phase, assuming a colony has already reached this stage upon appearance. It is not difficult to observe the initial exponential part of the colony growth (occurring before a colony reaches a radius of ∼100µm) by using commercially available photographic macro lenses (Fig. S6). However, one should be aware that defining appearance time with a linear regression is incorrect in this case as the exponential growth phase is actually observed and should be taken into account. Similarly, our estimation of appearance time from endpoint images assumes that colonies are in the linear growth phase.

As soon as the steady flow of nutrients sustaining colony growth decreases (timing is dependent on proximity of competing neighbors and the total number of colonies on the plate), the colony enters saturation phase, which impacts the linear appearance time estimation. Density may even affect the observed appearance time if colonies enter saturation phase before they are visible (Levin-Reisman *et al*., 2010). However, in the presented *S. aureus* dataset (including extremely dense plates), only a marginal correlation was observed between colony appearance time and density (SI Fig. 11). Colony growth is a complex biological process and different species on different growth substrates will respond differently to density (Chacón *et al*., 2018). For this reason, it is impossible to propose a universally valid method to take density into account. Nevertheless, the explored correction based on spatial metrics, which can easily be exported with ColTapp, might be used as a starting point to develop suitable correction methods for the investigated species.

We wrote the ColTapp application using the classical programming language MATLAB, and the code is designed as a modular shell that can host further image analysis methods that may meet specific needs, while benefiting from the easy to operate graphical interface. We enable the user to export all generated data, including radius, appearance time, growth rate, spatial metrics, and colony characteristics such as shape and color. We envision that more measurements can subsequently be added to facilitate the further study of intra- and inter-species colony interactions.

## Materials and methods

### Software

We programmed the ColTapp application with MATLAB 2020a (MathWorks). Preliminary code of the GUI and time-lapse radius tracking was proposed earlier (Vulin *et al*., 2018). All algorithm accuracy and computational efficiency tests were performed on a computer with Windows 10 OS, i7-8700K @4.70GHz, 32GB RAM.

We used external code acquired from the File Exchange server of MathWorks for certain functions. Namely, these were a Voronoi tessellation function by Sievers (2020), an image zoom and pan function by Cabrera (2020), multiple directory acquisition function by Cannell (2020), a smoothing function tolerant to missing values by Pittam (2020) and a function to fit a circle through three points by Malyuta (2020).

Additionally, we used ggplot2 (Wickham, 2016) within R and R Studio (R Core Team, 2018) as well as Inkscape for certain figures.

### Availability

The ColTapp MATLAB source code and executable (macOS/Windows) can be found at https://github.com/ColTapp under the GNU General Public License 3 and freely downloaded, together with a package of demo images and the code to trigger cameras with an Arduino board.

### Image acquisition

ColTapp does not rely on a specific image format or quality. Achieving homogenous lighting is undoubtedly critical for successful image analysis, but image acquisition itself can be operated using any kind of device. For correct spatial calibration, the camera should be positioned perpendicular to the plate. For time-lapse imaging, acquisition triggering at defined time intervals is necessary. Typical systems include scanners, dedicated applications and cameras or commercially available consumer-grade camera, provided they include a time-lapse mode or can be triggered by an external controller.

To generate the data presented in this paper, the time-lapse images were acquired either using a Canon EOS 1200D reflex camera triggered through an Arduino controller (Figure 1A) or Basler acA5472-17um 20MP cameras with Fujinon Objectives CF16ZA-1S 16mm/1.8M37.5×0.5 triggered with Basler’s pylon software suite (Version: 6.0.1). The plates were rested on a black background surface below the cameras and the whole system was set inside an incubator at 37° C. To prevent drying out of agar plates and airborne contamination, lids were kept on plates used for time-lapse imaging. Best image analysis quality was observed when special attention was brought to the illumination method: indirect, homogeneous white illumination was obtained by covering the incubator walls with white paper and using a LED based light source emitting minimal heat. Endpoint images were acquired manually with a Canon EOS 1200D reflex camera and lids were temporarily removed. Plates were placed in a custom-built box with light diffusing sides and a fixed plate- and camera-holder.

### Bacterial strains and Cultivation

*Staphylococcus aureus* strain Cowan I was grown in shaking test tubes containing tryptic soy broth (Difco) inoculated at an optical density of 0.2 with or without rifampicin (1 µg/ml). The control dataset was generated by plating appropriate dilutions of the non-treated cultures at mid-exponential phase. For the test dataset, the rifampicin exposed cultures were recovered after 24 hours, pelleted (10’000 g, 3 minutes), resuspended twice in phosphate buffered saline solution and then plated at appropriate dilutions.

Blood agar plates (Columbia, 5% sheep blood) were purchased at BioMérieux. Other strains and media used are described in the main text.

## Author details

**Julian Bär**

Department of Infectious Diseases and Hospital Epidemiology, University Hospital Zurich, University of Zurich, Zurich, Switzerland

**Contribution**: Writing of ColTapp code, performing experiments and analysis, writing of manuscript

**Competing Interests**: No competing interest declared

**Mathilde Boumasmoud**

Department of Infectious Diseases and Hospital Epidemiology, University Hospital Zurich, University of Zurich, Zurich, Switzerland

**Contribution**: Writing of ColTapp code, performing experiments and analysis, writing of manuscript

**Competing Interests**: No competing interest declared

**Roger Kouyos**

Department of Infectious Diseases and Hospital Epidemiology, University Hospital Zurich, University of Zurich, Zurich, Switzerland

Institute of Medical Virology, University of Zurich, Switzerland

**Contribution**: Conceiving and planning of project, reviewing the manuscript, supervision

**Competing Interests**: No competing interest declared

**Annelies S. Zinkernagel**

Department of Infectious Diseases and Hospital Epidemiology, University Hospital Zurich, University of Zurich, Zurich, Switzerland

**Contribution**: Conceiving and planning of project, reviewing the manuscript, supervision, acquisition of funding

**Competing Interests**: No competing interest declared

**Clément Vulin**

Department of Infectious Diseases and Hospital Epidemiology, University Hospital Zurich, University of Zurich, Zurich, Switzerland

Formerly: Institute of Biogeochemistry and Pollutant Dynamics, ETH Zurich, Zurich 8092, Switzerland Formerly: Department of Environmental Microbiology, Eawag, Dubendorf 8600, Switzerland **Contribution**: Conceiving and planning of project, Writing of ColTapp code, performing experiments and analysis, writing of manuscript

**Competing Interests**: No competing interest declared

## Acknowledgement

We thank Alejandra Rodriguez Verdugo, Anna-Karina Pitol and Florien Gorter for providing the pictures of plates with phage and bacteria. We thank Srikanth Mairpady Shambat for providing feedback on the manuscript.

