## Supplementary material for "ColTapp, an automated image analysis application for efficient microbial colony growth dynamics quantification": SI_text_and_figures

### Extended description of the implementation

#### 1 Graphical user interface

Global operating buttons to load images and save data are in the top left part of the graphical interface (Fig. S1-a) and information about currently operating functions and text feedback in the top right part (Fig. S1-b). Upon loading a folder containing images to analyze, the user needs to define the operating mode (*Time-lapse* (TL) mode or *Endpoint* (EP) mode), which cannot be changed for further analysis. Images in a folder are sorted with a natural sorting algorithm by name and assigned a frame number (from 1 to number of images in the folder). The currently active frame is visible at the center of the graphical user interface (Fig. S1-c). The user can switch between frames by moving the slider, typing the number of the desired frame (Fig. S1-d), navigating with the buttons on the sides of the image or pressing left/right arrow keys. The user can choose to display available data on the current frame in the *Options* menu, which also contains additional parameter configuration and gives access to other functorialities (Fig. S1-e). The main functionalities of ColTapp are grouped in three tabs on the left of the interface: *Detect*, *Main* and *Visualize* (Fig. S1, f g and h respectively). Two versions of the main panel are shown (Fig. S1-g) because the panel is dynamically adapted depending on the mode chosen by the user (Main-TL, Main-EP).

#### 2 Analysis setup

##### 2.1 Spatial calibration

The program proposes to define a pixel to µm spatial calibration factor. The user may either directly enter the factor if known, or define the diameter of a circle, typically the agar plate (automatically detected), or any reference line of known length on the image.

##### 2.2 Area of interest

Automatic colony detection can be narrowed to the defined boundaries of an area of interest (AOI). Users can select the plate as AOI or draw a custom polygon on the image. Using an AOI is suggested to reduce computational time and reduce false positive detection outside the boundaries of e.g. an agar plate. In *time-lapse* mode, both the spatial calibration factor and AOI are propagated from the active frame to the other frames. In *Endpoint* mode, these variables are only applied to the current image. This can then be manually propagated if all images in a given folder were acquired with the same setup in *Options*: *Apply calibration factor/AOI to all frames*.

### 3 Colony detection

#### 3.1 Color image conversion

ColTapp offers 16 methods to transform a color image into grayscale. Using one of the three RGB channels is computationally quick and generally precise enough, but in some special cases the user may want to use a different conversion as for example a transformation to other classical color spaces (e.g. CIELAB) and using one of the generated channels. The inbuilt MATLAB function rgb2gray which uses a weighted image conversion, retaining information of all three color channels, is also available.

#### 3.2 Algorithm parameters

A selection of parameters involved in circle detection and quality control can be tuned to potentially improve performance, although we tested the default parameters values on a selection of images and reported high accuracy. The range of expected radius is an exception and we suggest to always set it as narrow as possible. This range greatly affects computational speed and accuracy of colony detection. The range boundaries (expected minimal and maximal radius in pixels) can be either derived automatically from the smallest and biggest circle drawn on the currently displayed image with the function *Define radius range* or directly specified within the *Options* dialog. The following other parameters can be modified: the relative size of $img_{crop}$, minimal distance from $img_{crop}$ boundaries, a bias towards foreground classification as well as the minimal proportion needed to be classified as foreground within a circle. Maximal proportion of a circle overlapping with one other circle as well as minimal distance and radius difference are modifiable. Additionally, the user can set maximal total area of a circle overlapping with any other circle and define the final minimal distance between any circle.

#### 3.3 Results corrections

Addition of non-detected colonies and false positive removal is possible through simple mouse-guided operations. Additionally, polygons can be drawn on the image to clear out entire zones from found circles. In *Time-lapse* mode, it is important for downstream analysis to check if none of the detected circles is composed from two (or more) colonies which merged during the time-lapse imaging. These should be removed and replaced with the appropriate number of circles with corresponding centers. This control step is needed to be done manually.

### 4 Colony characteristics

#### 4.1 Shape and color metrics

ColTapp can extract basic shape and color parameters from the colonies on each frame for further analysis, such as species identification or observation of mutation induced phenotypes linked with altered growth on agar plates. Color of the colonies can be extracted from the entire colony or from the center (within a 5-pixel radius), and texture is calculated either as standard deviation of the color or image entropy (Simunovic *et al.*, 2016). The length of the perimeter of colonies as well as the standard deviation on the average radius are metrics useful to categorize colonies based on the shape of the border. We also propose to export the mean color of the halo around colonies as this could be useful for colorimetric assays (Palkova *et al.*, 2002) or quantification of hemolysis capacities of the colony.

##### 4.2 Spatial metrics

For post-processing steps such as density correction or the study of colonies interactions, the user can export typical spatial metrics calculations. Interactions between neighboring colonies are mostly occurring through the diffusion of small molecules through the agar plate matrix. The diffusion in such 3D gels can sometimes be approximated to be two-dimensional, and basic solutions of the diffusion equations will give interactions based on the distance D between colonies either as $\frac{1}{D}$ or $\frac{1}{D^2}$ depending on how the diffusion equation is solved. In their study to estimate parameters influencing colony size on plates with multiple colonies, Chacón *et al.* (2018) assumed that interactions from neighboring colonies are additive, resulting in interaction terms defined by $\sum\frac{1}{D}$ and $\sum\frac{1}{D^2}$ . We implemented calculation of these metrics within ColTapp. We hypothesize that not only the location but also the size of colonies surrounding the focal colony is likely to play a role in the interaction magnitude. Thus, we calculate an additional metric: the sum of angular diameters, calculated as $AD = \sum 2\arctan(\frac{R_i}{D_i})$, where $R_i$ and $D_i$ are the radius and distance from the focal colony of each neighboring colony. This metric, unexplored in the context of colony growth, is biologically plausible since it assumes that distant large colonies (that have consumed large amount of resources) have a similar effect as small but closer colonies, consuming resources closer to the focal colony. A user may define distance cutoffs to limit the interaction terms to colonies within a maximal distance to the focal colony for further calculations.

Finally, Chacón *et al.* (2018) proposed that the amount of nutrients available to each colony on the agar plate can be approximated with Voronoi cell areas, obtained by tracing perpendicular bisector lines between each pair of neighboring colonies (Okabe *et al.*, 2009). They showed that the final size colonies attain when agar plate carrying capacity is reached can be predicted by the colonies Voronoi cell areas. ColTapp computes these areas by performing a Voronoi tessellation within user-specified boundaries (Sievers, 2020).

### 5 TL-Preprocessing steps

#### 5.1 Drift correction

Slight drift can occur in time-lapse setups during imaging. Therefore, ColTapp incorporates an image registration algorithm which performs a 2-D rigid translation at subpixels resolution ($1/\kappa$), where the factor $\kappa$ (adjustable as *RegistrationFactor* in *Options*) can be specified by the user. The algorithm computes the up-sampled cross-correlation with a discrete Fourier transform (Guizar, 2020). The drift correction is done on a user-defined small area of the image by generating a translation vector in respect to the reference frame. Concretely, the translation is not applied to the images themselves, but to the colonies' centers position.

#### 5.2 Colony centers correction

The centering of colony circles is crucial to track radii over time, especially at early timepoints when colonies are small. ColTapp offers an automatic center correction, by tentatively detecting circles on sub-images cropped from an early frame based on the location and radii of the colonies detected on the reference frame. These newly detected circles' center coordinates are kept as colony center, unless they are farther away than a user-defined threshold from the original coordinates or if they are set to the center coordinates of another colony in close proximity. In these cases, the correction is skipped, and the colonies are added to a list for subsequent manual center correction. If no circle is detected at all because the colony is not yet visible at that timepoint, the same process is repeated 10 frames later. The user is able to monitor this process visually.

Alternatively, the user may choose to execute this task manually, by clicking the correct center on the sub-images sequentially displayed.

### 6 Tracking radii over time

#### 6.1 Overlapping colonies

Neighboring colonies can be visible as blurry regions in the top of kymographs, hindering creation of clean binary images (Fig. 5B). To reduce this phenomenon, before the kymograph creation process, overlapping colonies are automatically detected and the ranges of angles corresponding to adjacent colonies are discarded from the polar transformed intensity data. If more than 90% of all angles are discarded because of overlap, the exclusion of angles is omitted completely, to avoid reducing the available data too much. The overlap detection functionality can be deactivated or tuned with *Scale radius for overlap* (accessible in *Options*). This scaling factor is multiplied to the radius of the focal colony from which center neighboring colonies are tested for overlap: by increasing it, a user may choose to discard ranges of angles corresponding not only to overlapping colonies but also very close colonies. Note that this increase might lead to high proportions of angles to be discarded. Decreasing the scaling factor leads to reduced ranges of excluded angles. This might be useful in densely populated plates to still achieve some overlap exclusion to increase quality of kymographs at earlier timepoints.

##### 6.2 Radial growth curve correction

ColTapp has an inbuilt function to automatically detect radial growth curves with probable errors based on the number of local maxima, size of radius differences from frame to frame, monotonicity, and number of frames without a successful radius determination. These (or any) radial growth curves can be manually corrected with a dedicated tool (Fig. S3) which allows the user to switch for each colony individually between Global thresholding and Edge detection method, and adjust any parameter of the two methods to derive best parameter combinations to derive the radial growth curve from the kymograph.

#### 7 Reference growth parameters definition

The user can define the reference radial growth rate and appearance time either by providing known values manually, or by directing ColTapp to a folder containing a control experiment monitored by time-lapse and already analyzed, to extract these parameters automatically. In this case, ColTapp obtains the mean of appearance time and radial growth rates of the control experiment's colonies. Users concerned by the presence of outliers in the growth control experiment may change from mean to median (or any user-defined quantile) in the *Options*. These reference experiments can be used for further calibration.

#### 8 Sequence of endpoint images

##### 8.1 Delayed colony detection

Some late appearing colonies may not be observed at the optimal time for colony size observation (e.g. 24h). In that case, we suggest capturing image(s) at later timepoint(s) (e.g. 48h). ColTapp allows the user to overlay images taken from the same plate at different times to ease detection of late growing colonies which did not yet appear at an earlier timepoint. ColTapp assumes that a folder under analysis contains multiple images from the same timepoint and requires the user to navigate to a folder containing the same set of images with matching order, acquired at another timepoint.

To achieve similar image orientation at both timepoints, the user may place a mark on the edge of the plate as reference. Users can manually align images which have not been captured in the exact same orientation by clicking at two matching positions on the images of the two timepoints.

The inbuilt MATLAB function *fitgeotrans* is used to fit a geometric transformation to the pairs of points with a nonreflective similarity to estimate rotation and translation of the two images to align. If delayed colonies are revealed at this point, the user has the option to add not yet visible colonies to the earlier timepoint, which are defined as colonies with a radius equal to zero.

##### 8.2 Sequence of endpoint images

The user can link corresponding folders of multiple image sets taken at different timepoints to create timeseries data without the need for a time-lapse imaging setup. Colonies on the images in different folders are matched based on the minimal distance of centers between colonies. Mismatches can be corrected by aligning the images at different timepoints by fitting a geometric transformation to user-defined pairs of points with a nonreflective similarity to estimate rotation and translation of the two images to align. It is necessary that the number of detected colonies is the same in all folders.

#### 9 Visualization tab

Data previewing is possible through functions accessible within the *Visualization* tab with slightly differing options for *time-lapse* and *endpoint* mode. Size visualizations can either be done in pixel or micrometer units (*Options*). Radius distributions can be displayed for a user-defined frame of a time-lapse or a user-defined image of a set of endpoint images while also giving the option to combine all images of a given folder into one distribution. In *time-lapse* mode, the radial growth curves can be visualized. Time can be set as either as frames or hours (*Options*). Appearance and growth rate distribution visualizations are also implemented. The number of bins used for distributions can be set within *Options*.

#### 10 Options

Parameters of implemented functions can be set by the user within the *Options* window (Fig S2). The *Options* window is divided in four tabs (*Global*, *Detect*, *Main* (*TL* or *EP*) and *Visualize*). We describe some of the most important parameters and functions here.

The *Global* tab allows to set the method for converting a colored image into a grayscale version, activate direct data visualization options on the image (e.g. colony circles and Voronoi edges), set averaging strategy for reference growth data extraction and to disable autosaving.

The *Detect* tab contains options to set image binarization method (Adaptive (local), Otsu (global) or none), invert the image for dark colonies on bright background, choose colony detection method (Regionprops based circle detection or direct circle detection) and modify any of the parameters of the colony detection algorithm.

The *Main-TL* tab allows to tune parameters related to radial growth curve creation and appearance time determination. For example, definition of the default kymograph processing method, time interval between frames, *RegistrationFactor*, scaling of radius for overlap exclusion as well as $R_{thresh}$ and $Fr_{lin}$ are possible. Additional functions are subset radii tracking, manual timepoint specific radius correction, curve removal and interactive scaling of the radii defining the size of $img_{crop}$.

The *Main-EP* tab contains small functions to assign the AOI and spatial calibration factor of the current frame to all frames of the loaded folder and remove the linked folders.

Finally, the *Visualize* tab allows to change details of the data previewing functions. Smoothing factor for radial growth curves, units to use for plotting (pixel or micrometer and frame or hour), number of bins used for distributions and titles can be set here.

### 11 Data export

The generated data is automatically saved in the same folder as the images are stored in the native MATLAB format (.mat files) when running the application. For export to another software, ColTapp provides a data export functionality. Through the export menu, user can save comma separated values files (delimiter can be changed) of position, radius, fitted appearance time, as well as several calculated density measures and simple shape and color parameters of each colony.

Most variables are exported as a three-column table, containing the frame number, the colony number and the exported data of interest, grouped with similar variables. In *time-lapse* mode, the radius of colonies is exported as a matrix of time by colony to facilitate extraction of growth curves.

The user can choose to export relevant variables either as pixel values or as µm values, provided the spatial calibration factor was set on all frames. Additional data as for example the spatial calibration factor, used color to grayscale transformation method as well as plate center and radius (if specified) can be exported in a separate metadata csv file.

### Figures

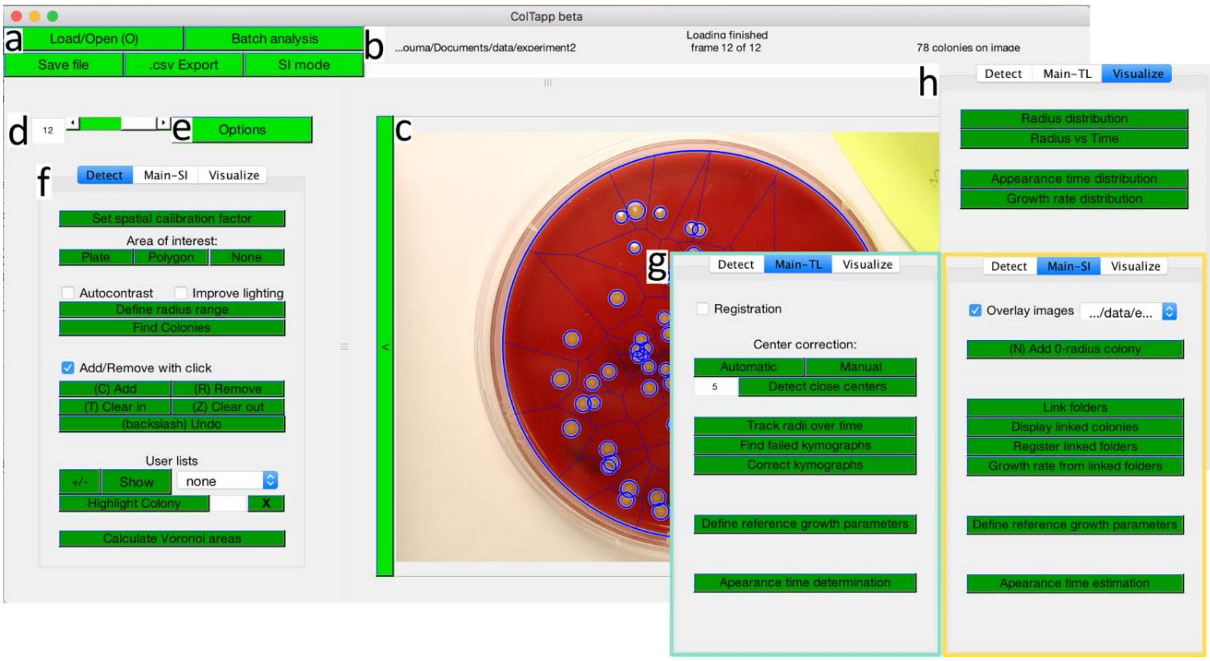

**Supplementary Figure 1:** The ColTapp graphical user interface, described in SI text 1.

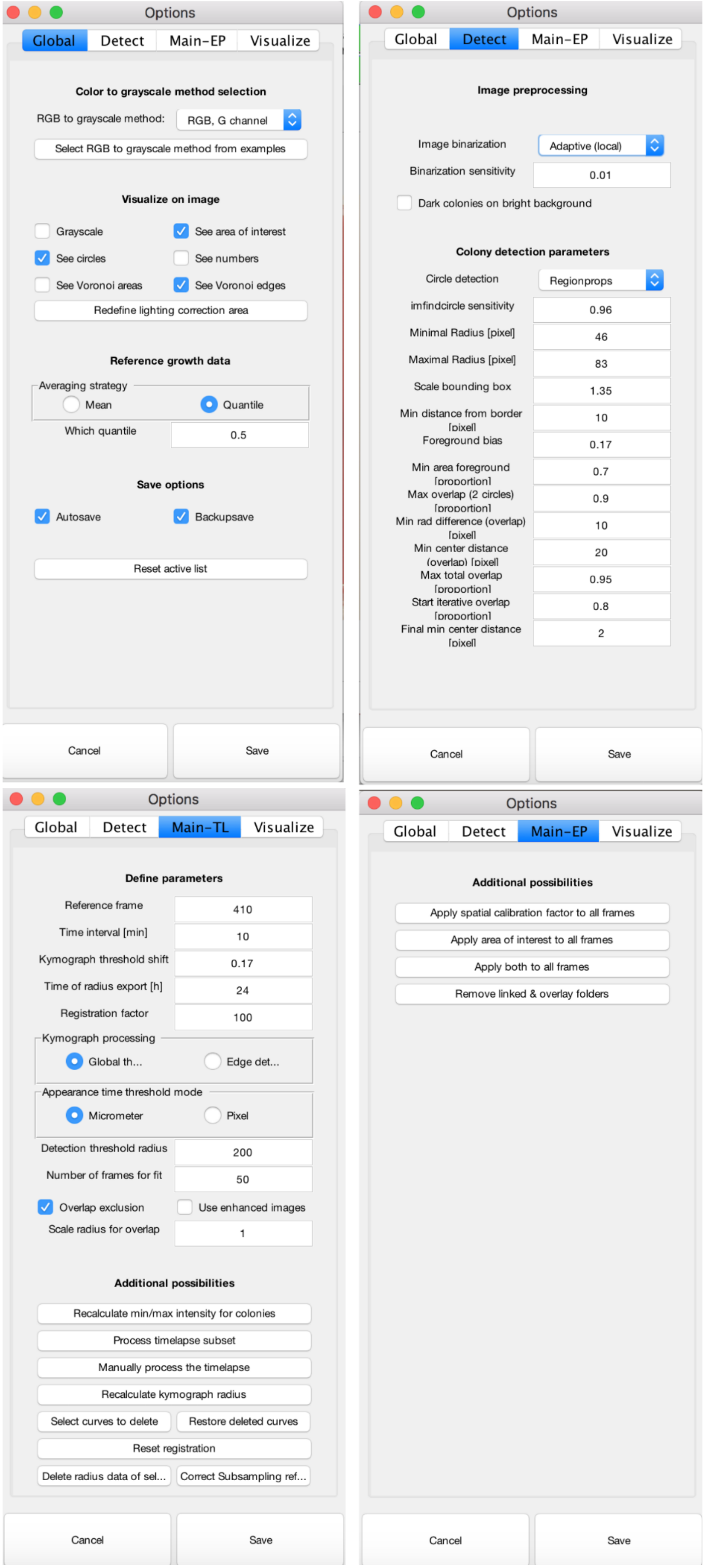

**Supplementary Figure 2:** The ColTapp options window, described in SI text 10.

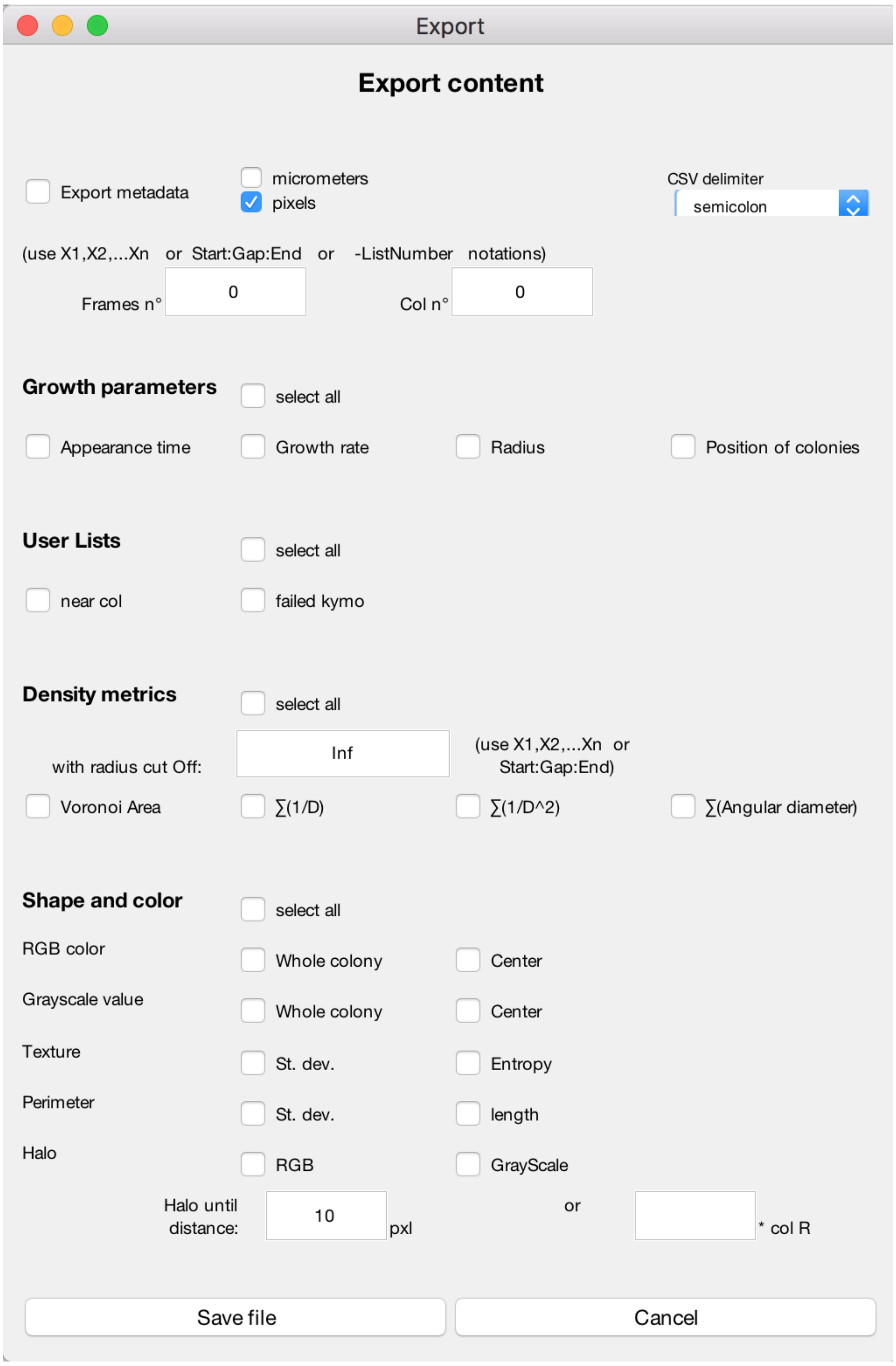

**Supplementary Figure 3:** The ColTapp export window. The export functionality is described in SI text 11.

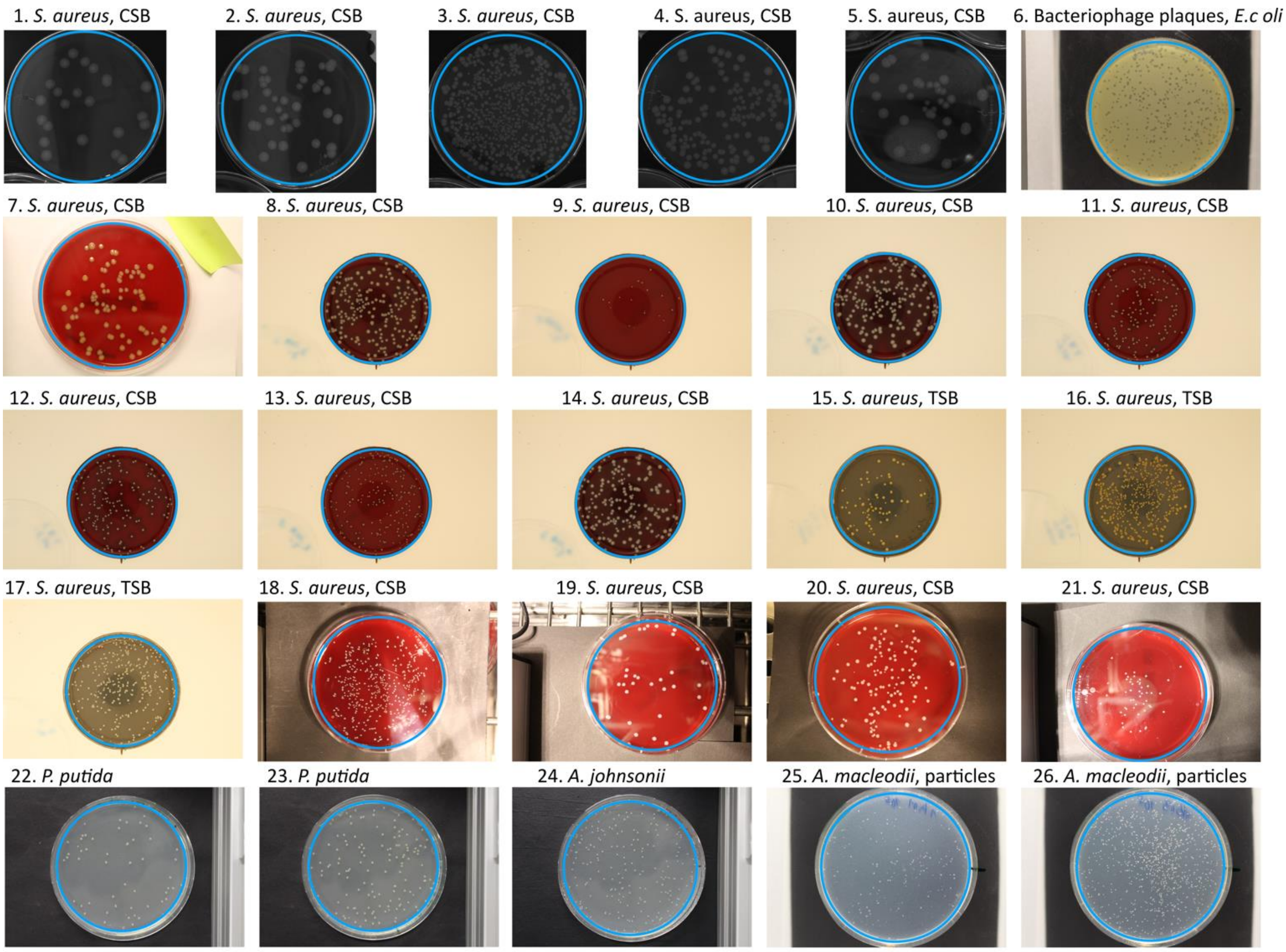

**Supplementary Figure 4:** The 26 images used for assessment of computational time and accuracy of the colony detection algorithm (Table S1). The area of interest was set to the displayed blue circle on each image. CSB: Columbia sheep blood, TSB: tryptic soy broth.

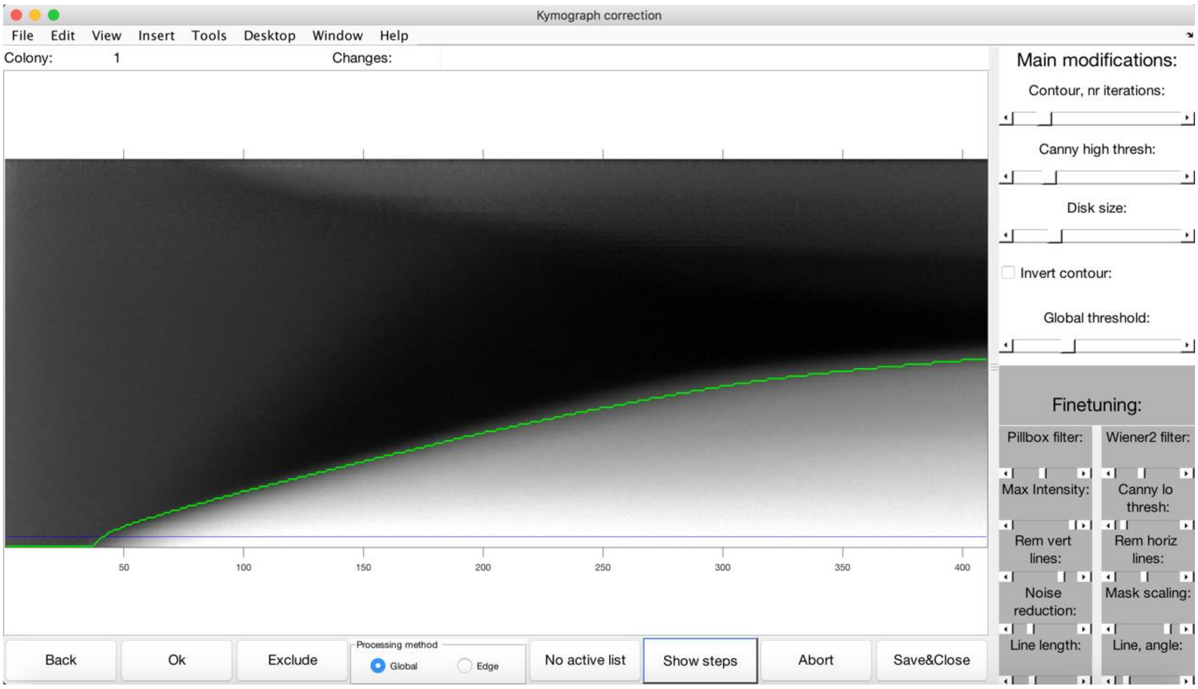

**Supplementary Figure 5:** The ColTapp kymograph correction window. The user can visualize kymographs and the derived radial growth curve (shown here in green) sequentially and correct the growth curve manually. If the Global thresholding method does not perform well, a user may choose to change to the Edge detection method. All parameters for both methods are tweakable with the sliders in the panel on the right side. Additionally, buttons to exclude the currently displayed colony, adding it to a list and display all steps of the Edge detection method in a popup window for better determination of the source of error are available.

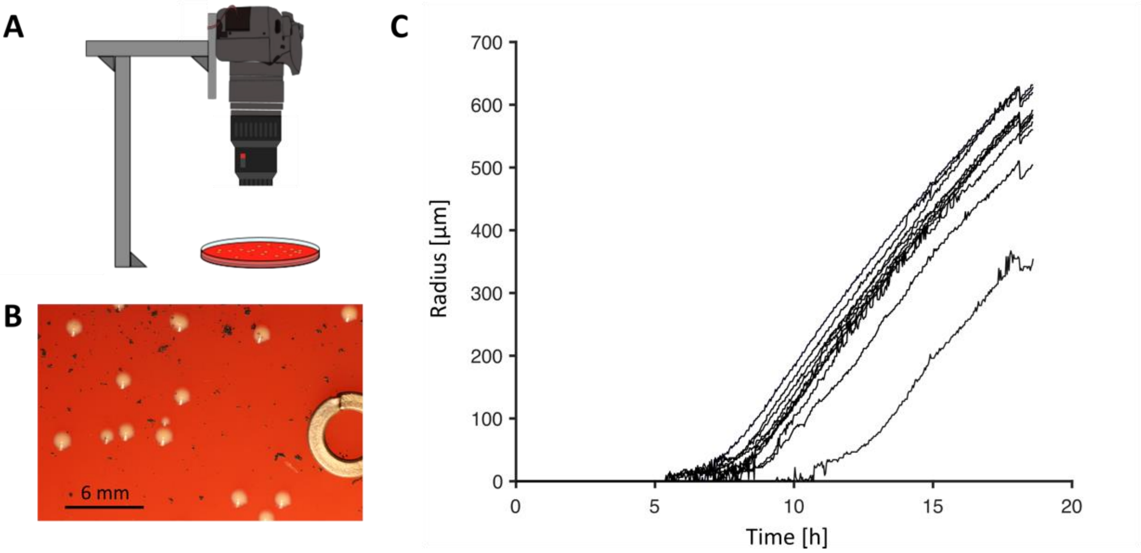

**Supplementary Figure 6:** Close-up colony growth. **A.** The zoom factor of the images can easily be increased to a level were the exponential phase of bacterial colony growth can be observed, by mounting commercially available macro lenses to a camera. **B.** As an example, *S. aureus* bacterial colonies are grown on a blood plate. Some particles of activated charcoal are visible on the plate to help with focusing when experiment is setup, and a metal ring on the right of the image serves as a distance reference. **C.** With this setup, the distance resolution can be brought below 50 µm, and a short exponential phase is observed when colonies grow, before they reach ~100 µm. In this case, the appearance time cannot be estimated using a simple linear regression and calculations will need to be adapted.

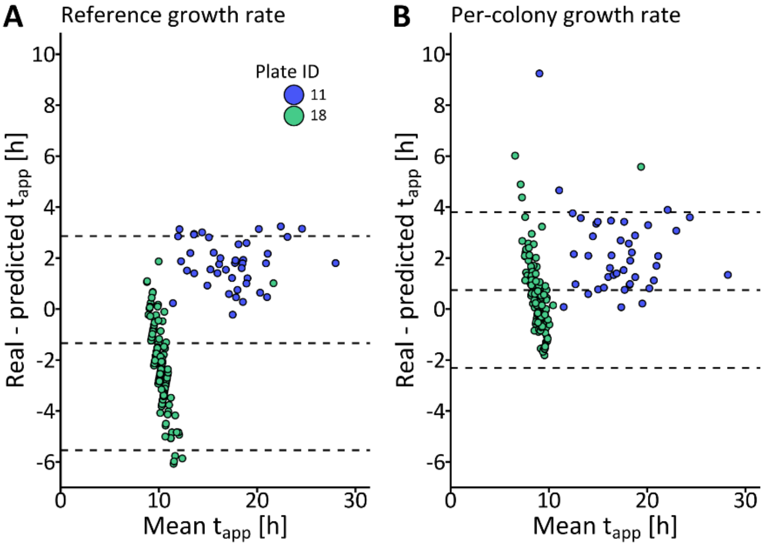

**Supplementary Figure 7**: Bland-Altman plots of appearance time estimation from endpoint images using **A.** a reference growth rate of 65 µm/h and **B.** per-colony estimated growth rates from a sequence of endpoint images.

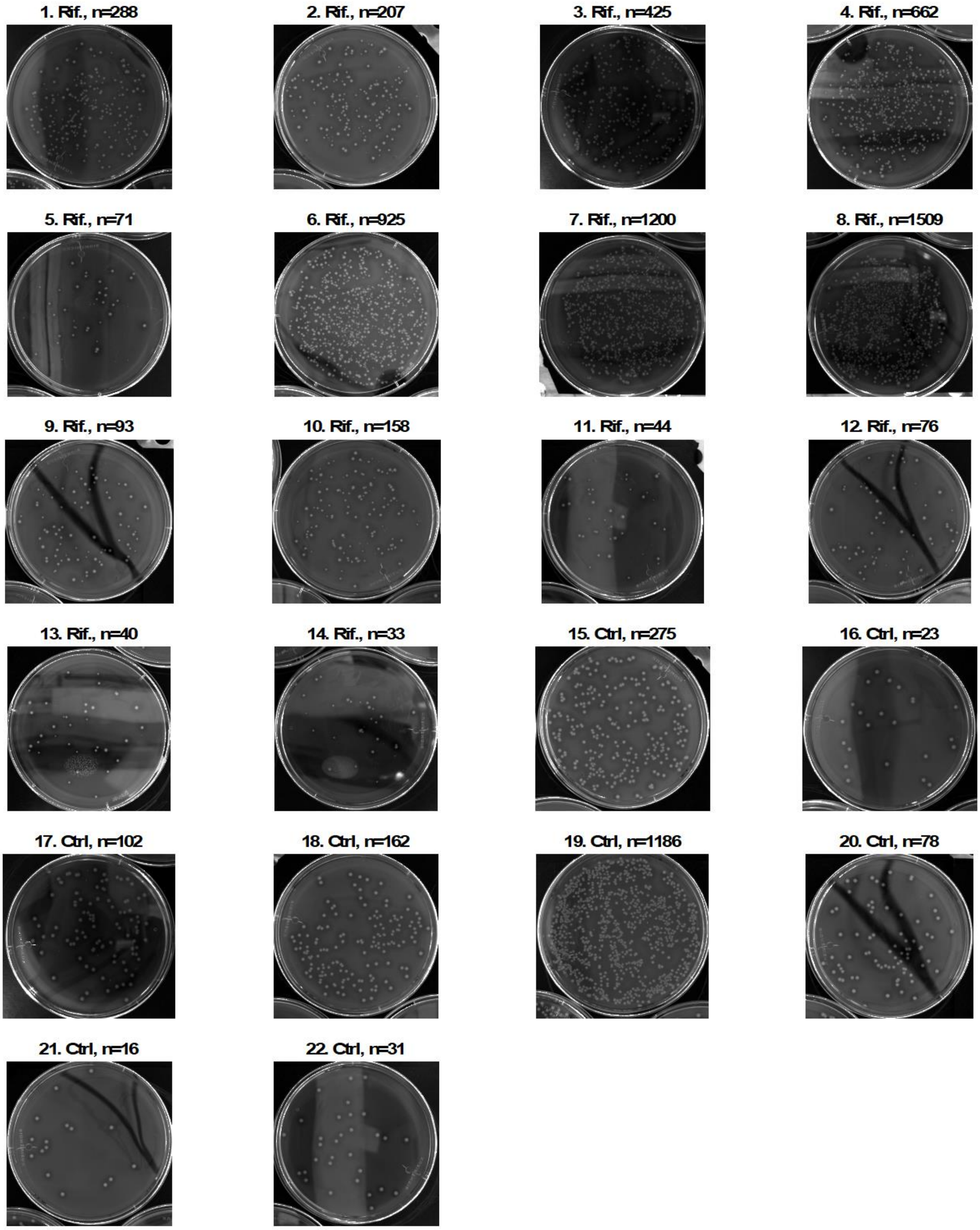

**Supplementary Figure 8:** Images of the demonstration dataset plates at 24 h. Ctrl: Control dataset, Rif: Rifampicin treated dataset, n: number of colonies on plate

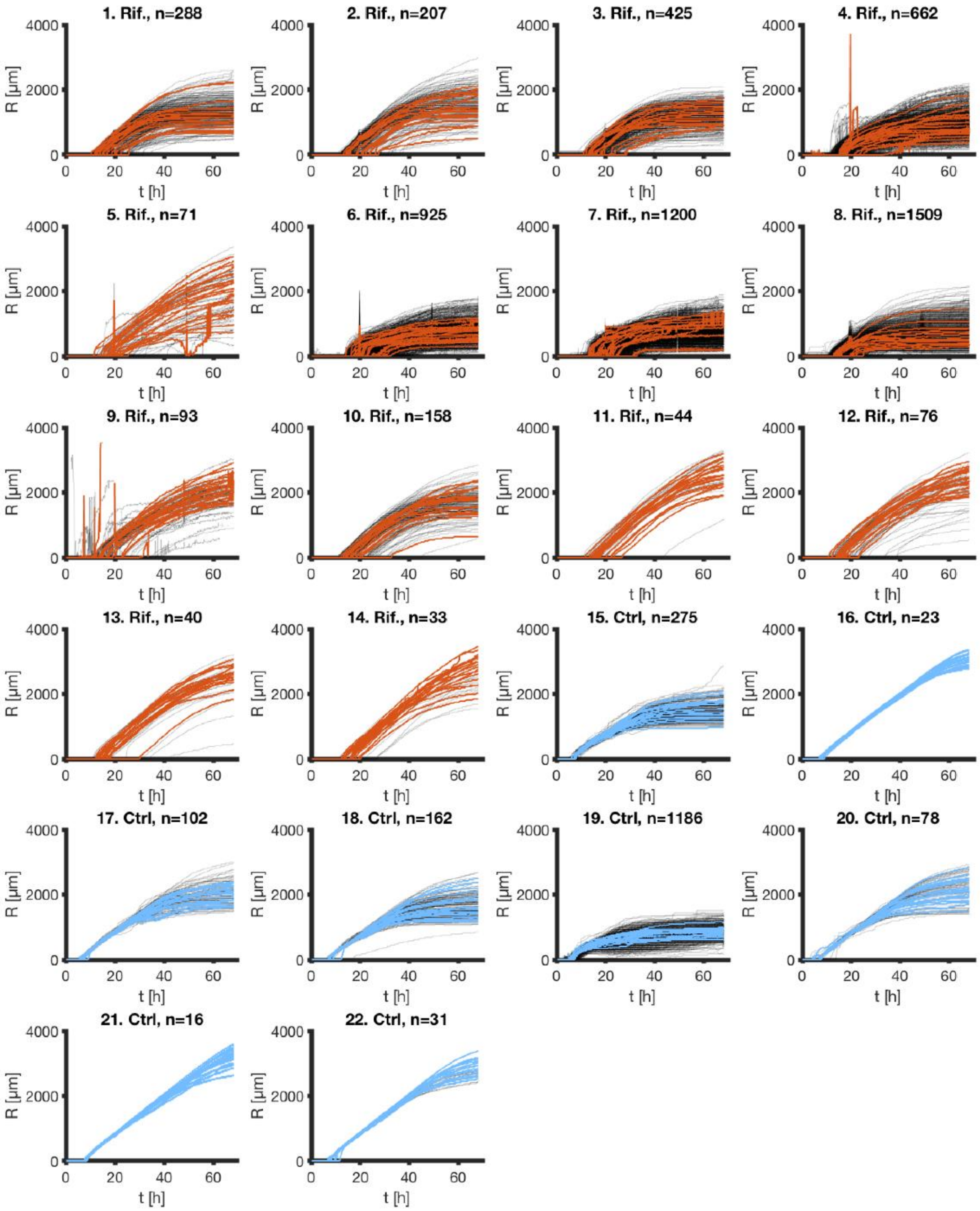

**Supplementary Figure 9:** Growth curves obtained for the control (blue) and rifampicin treated (orange) dataset. All curves are displayed in black and 20 random growth curves are colored for each plate to illustrate the general dynamics.

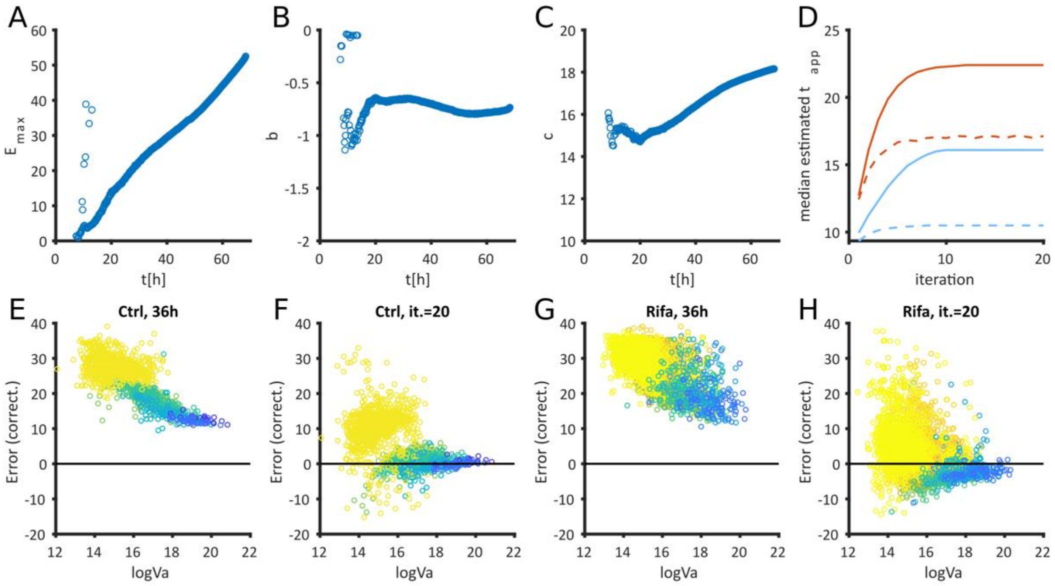

**Supplementary Figure 10:** Panels **A, B** and **C** show the values of the parameters obtained by fitting the control dataset (panel E) at different timepoints (see also SI movie 3). The maximal error $E_{max}$ linearly increased through time while the predicted values of the parameters b and c are fairly consistent through time apart from early timepoints. **D**. Representation of the median predicted $t_{app}$ on each iteration of the correction shows that the predictions stabilize in fewer than 10 iterations. Panel **E** and **G** show the uncorrected error of plates at late time point (36h) for the control and the rifampicin dataset, respectively. Panel **F** and **H** are the corrections obtained after 20 iterations.

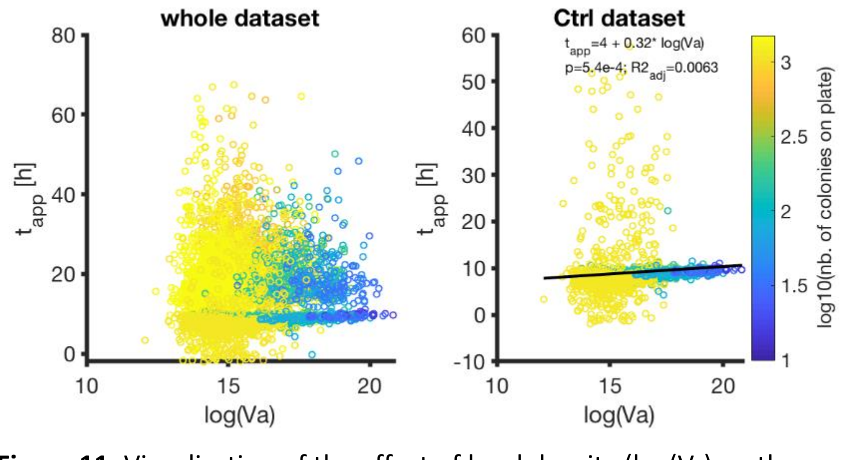

**Supplementary Figure 11:** Visualization of the effect of local density ($\log(V_a)$ on the appearance time, either on the whole dataset (**left**) or solely on the control dataset (**right**). We didn't notice an important effect of density on $t_{app}$, and we attribute the observed slope to noise in the dense plate data.

### Tables

| Image Nr. | Color space | Species | Total time [s] | Time per circle [s] | N initial circles | N false positive | N false negative | N corrected | False positve rate [%] | False negative rate [%] | False positive in clusters | Average radius [px] | SD radius [pixel] | Min. expected radius [pixel] | Max. expected radius [pixel] |
|---|---|---|---|---|---|---|---|---|---|---|---|---|---|---|---|
| 1 | Grayscale | S.aureus | 2.22 | 0.1009 | 22 | 0 | 1 | 23 | 0.00 | 4.35 | no | 62.91 | 3.94 | 45 | 77 |
| 2 | Grayscale | S.aureus | 2.38 | 0.0553 | 43 | 0 | 1 | 44 | 0.00 | 2.27 | no | 52.66 | 9.03 | 36 | 64 |
| 3 | Grayscale | S.aureus | 9.59 | 0.0125 | 767 | 7 | 165 | 760 | 0.91 | 17.84 | no | 17.87 | 5.94 | 8 | 39 |
| 4 | Grayscale | S.aureus | 3.51 | 0.0239 | 147 | 0 | 5 | 152 | 0.00 | 3.29 | no | 34.53 | 7.54 | 22 | 63 |
| 5 | Grayscale | S.aureus | 2.97 | 0.0990 | 30 | 1 | 3 | 32 | 3.33 | 9.38 | no | 54.53 | 11.28 | 26 | 89 |
| 6 | RGB | Phage plaques | 12.76 | 0.0264 | 484 | 11 | 33 | 506 | 2.27 | 6.52 | no | 18.66 | 6.61 | 8 | 35 |
| 7 | RGB | S.aureus | 9.2 | 0.1150 | 80 | 3 | 1 | 78 | 3.75 | 1.28 | no | 54.10 | 8.76 | 35 | 76 |
| 8 | RGB | S.aureus | 6.86 | 0.0306 | 224 | 2 | 2 | 224 | 0.89 | 0.89 | no | 33.23 | 5.13 | 10 | 27 |
| 9 | RGB | S.aureus | 3.7 | 0.0822 | 45 | 25 | 0 | 20 | 55.56 | 0.00 | yes | 14.15 | 3.42 | 6 | 29 |
| 10 | RGB | S.aureus | 7.16 | 0.0405 | 177 | 0 | 5 | 182 | 0.00 | 2.75 | no | 38.59 | 5.42 | 26 | 69 |
| 11 | RGB | S.aureus | 4.94 | 0.0270 | 183 | 25 | 2 | 160 | 13.66 | 1.25 | yes | 20.72 | 2.54 | 9 | 31 |
| 12 | RGB | S.aureus | 4.86 | 0.0291 | 167 | 14 | 5 | 158 | 8.38 | 3.16 | yes | 20.02 | 2.25 | 11 | 32 |
| 13 | RGB | S.aureus | 5.35 | 0.0217 | 247 | 25 | 7 | 229 | 10.12 | 3.06 | yes | 14.58 | 3.17 | 7 | 32 |
| 14 | RGB | S.aureus | 7.6 | 0.0396 | 192 | 0 | 5 | 197 | 0.00 | 2.54 | no | 35.43 | 7.03 | 14 | 53 |
| 15 | RGB | S.aureus | 3.91 | 0.0455 | 86 | 13 | 1 | 74 | 15.12 | 1.35 | yes | 25.21 | 6.17 | 8 | 46 |
| 16 | RGB | S.aureus | 7.67 | 0.0161 | 476 | 5 | 21 | 492 | 1.05 | 4.27 | no | 18.56 | 4.86 | 8 | 38 |
| 17 | RGB | S.aureus | 6.6 | 0.0201 | 329 | 13 | 66 | 382 | 3.95 | 17.28 | yes | 17.73 | 3.11 | 8 | 27 |
| 18 | RGB | S.epidermidis | 1.94 | 0.0040 | 486 | 46 | 36 | 476 | 9.47 | 7.56 | yes | 16.46 | 2.74 | 12 | 31 |
| 19 | RGB | S.epidermidis | 6.81 | 0.0987 | 69 | 33 | 2 | 38 | 47.83 | 5.26 | yes | 42.10 | 2.70 | 53 | 56 |
| 20 | RGB | S.epidermidis | 9.34 | 0.0580 | 161 | 25 | 3 | 139 | 15.53 | 2.16 | yes | 31.79 | 7.62 | 11 | 57 |
| 21 | RGB | S.epidermidis | 5.97 | 0.0905 | 66 | 17 | 1 | 50 | 25.76 | 2.00 | yes | 21.37 | 1.55 | 17 | 36 |
| 22 | RGB | P.putida | 5.27 | 0.1033 | 51 | 0 | 3 | 54 | 0.00 | 5.56 | no | 21.62 | 2.06 | 19 | 33 |
| 23 | RGB | P.putida | 6.26 | 0.0467 | 134 | 1 | 12 | 145 | 0.75 | 8.28 | no | 21.42 | 1.76 | 16 | 33 |
| 24 | RGB | A.johnsonii | 7.08 | 0.0337 | 210 | 1 | 29 | 238 | 0.48 | 12.18 | no | 15.16 | 1.12 | 12 | 23 |
| 25 | RGB | A.macleodii | 7.04 | 0.0414 | 170 | 35 | 8 | 143 | 20.59 | 5.59 | no | 14.28 | 2.49 | 12 | 25 |
| 26 | RGB | A.macleodii | 16.84 | 0.0186 | 906 | 19 | 110 | 997 | 2.10 | 11.03 | no | 12.33 | 1.29 | 11 | 22 |
| | | **Average:** | 6.455 | 0.0492 | 228.9231 | 12.34615 | 20.26923 | 230.5 | 9.29 | 5.43 | - | 28.08 | 4.60 | 17.30769 | 43.96154 |
| | | **SD:** | 3.31762 | 0.0333 | 224.6649 | 13.20891 | 38.50564 | 240.7313 | 14.41 | 4.77 | - | 14.70 | 2.75 | 12.46682 | 19.06301 |

**Supplementary Table 1: Colony detection algorithm performance** (see caption below)

| Plate ID | N colonies | N frames | Total time [s] | Time per colony and frame [s] | Average radius [pixel] | N detected as failed | N false negative | N true negative | False negative rate [%] | N tn, need manual correction | N false positive | False detection rate [%] | N fp, need manual correction | N total false | Rate of failed, global [%] | N total need manual correction | Rate of manual correction needed [%] |
|---|---|---|---|---|---|---|---|---|---|---|---|---|---|---|---|---|---|
| 1 | 288 | 423 | 4049.44 | 0.0332 | 27.7263 | 50 | 3 | 47 | 6.00 | 16 | 9 | 84.75 | 1 | 56 | 19.44 | 17 | 5.90 |
| 2 | 207 | 423 | 3403.52 | 0.0389 | 31.2876 | 30 | 5 | 25 | 16.67 | 6 | 8 | 78.95 | 2 | 33 | 15.94 | 8 | 3.86 |
| 3 | 425 | 423 | 8248.11 | 0.0459 | 35.0090 | 119 | 2 | 117 | 1.68 | 23 | 14 | 89.47 | 1 | 131 | 30.82 | 24 | 5.65 |
| 11 | 44 | 423 | 1517.18 | 0.0815 | 51.4680 | 2 | 1 | 1 | 50.00 | 0 | 1 | 66.67 | 0 | 2 | 4.55 | 0 | 0.00 |
| 12 | 76 | 423 | 1938.10 | 0.0603 | 42.7522 | 12 | 3 | 9 | 25.00 | 3 | 2 | 85.71 | 2 | 11 | 14.47 | 5 | 6.58 |
| 13 | 40 | 423 | 1596.20 | 0.0943 | 58.1285 | 7 | 1 | 6 | 14.29 | 0 | 1 | 87.50 | 0 | 7 | 17.50 | 0 | 0.00 |
| 14 | 33 | 423 | 1181.89 | 0.0847 | 54.8332 | 4 | 0 | 4 | 0.00 | 1 | 0 | 100.00 | 0 | 4 | 12.12 | 1 | 3.03 |
| 15 | 275 | 410 | 9261.21 | 0.0821 | 33.0484 | 124 | 0 | 124 | 0.00 | 6 | 29 | 81.05 | 1 | 153 | 55.64 | 7 | 2.55 |
| 16 | 23 | 410 | 1792.52 | 0.1901 | 56.9671 | 3 | 0 | 3 | 0.00 | 0 | 1 | 75.00 | 0 | 4 | 17.39 | 0 | 0.00 |
| **Average:** | 156.78 | 420.11 | 3665.35 | 0.079 | 43.47 | 39.00 | 1.67 | 37.33 | 12.63 | 6.11 | 7.22 | 83.23 | 0.78 | 44.56 | 20.88 | 6.89 | 3.06 |
| **SD:** | 146.53 | 5.73 | 3043.15 | 0.047 | 12.08 | 49.33 | 1.73 | 49.37 | 16.62 | 8.15 | 9.48 | 9.43 | 0.83 | 58.24 | 14.75 | 8.49 | 2.65 |

**Supplementary Table 2: Track radius algorithm performance** (see caption below)

**Supplementary Table 1: Colony detection algorithm performance**. We tested the accuracy and computational efficiency of the colony detection algorithm with a set of 26 images (Fig S4). Detailed results for each image are displayed here. Average and standard deviation (SD) of each measurement are shown at the bottom of the table. We recorded number of detected circles by the algorithm, as well as the number of false positive of these and number of missed (false negative). We computed false negative and false positive rates with these numbers. Most of the images with high false positive rate had the false negative circles detected in large clusters, which is indicated on the table. Additionally, we measured computational time of the detection algorithm and report computational time per circle as well. Finally, average size of colonies in pixels and the range of expected radius is shown.

**Supplementary Table 2: Track radius algorithm performance**. We tested the accuracy and computational efficiency of the track radius algorithm with a subset of the dataset created for the density correction validation. Plate ID in the table are referring to ID numbers on Fig. S9 and S9. Detailed results for each time-lapse are displayed here. Average and standard deviation (SD) of each measurement are shown at the bottom of the table. For testing, we set the default growth curve determination method to Global thresholding. We report here the number of colonies and frames of each time-lapse as well as the number of radial growth curves automatically identified as failed after applying the Global thresholding method, the number of growth curves falsely categorized as failed (including rate), and the number of the identified growth curves which needed more manual correction than a simple switch to the Edge detection method. Corresponding rates. Next, we manually assessed all growth curves not classified as failed and record number of falsely categorized as correct (including false detection rate). Of these manually identified, we assessed if more manual correction than a switch to Edge detection method was necessary. The combined number of automatically and manually identified growth curves is reported here and the rate of failure of the Global thresholding method is given. Overall, only few growth curves need manual correction after switching to the Edge detection method. Additionally, we report the total computational time and the time per colony and frame as these two factors are proportional to required time. Finally, Average colony size at the last frame of the time-lapse is indicated.

### Movies

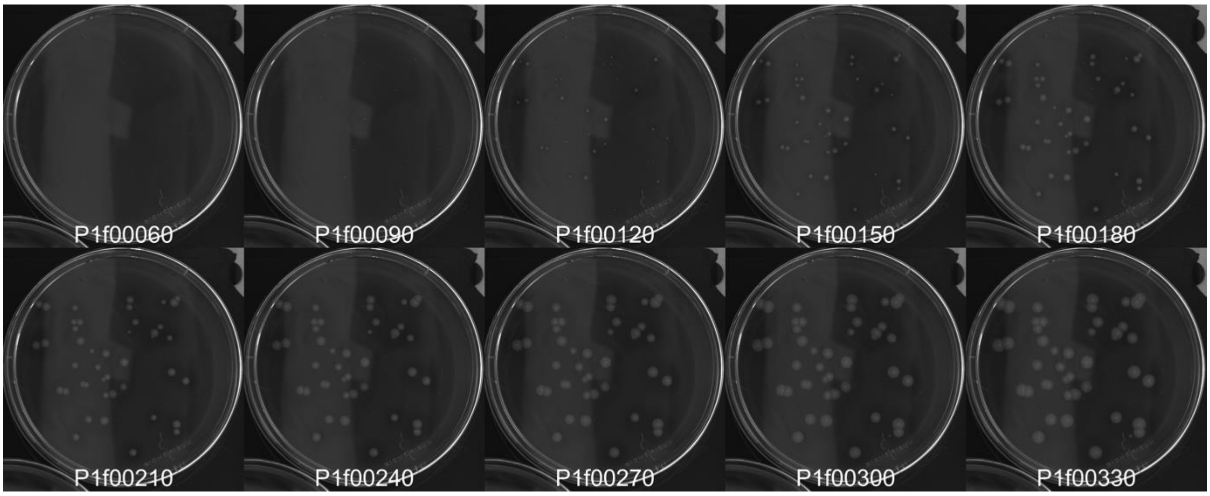

**Supplementary Movie 1:** Representative images of the SI movie 1 (one image every 5h, starting at 10h) that show a time-lapse of colony growth. The time-lapse movie is generated from the images of plate 11 (Fig. S9, Fig. S10). It includes 423 frames acquired in 10 min intervals.

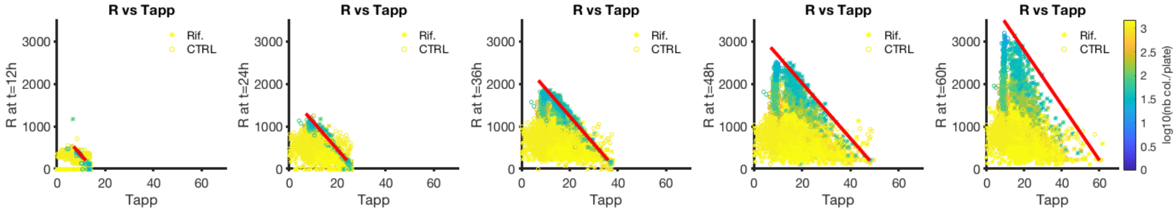

**Supplementary Movie 2:** This movie merges data of Fig 7B and D (full dataset) and shows the evolution of the correlation of radius of colonies and their appearance time. The red line represents the radius that could be obtained assuming the maximal growth rate, and deviations from this radius means that colonies grow slower than this maximal growth rate.

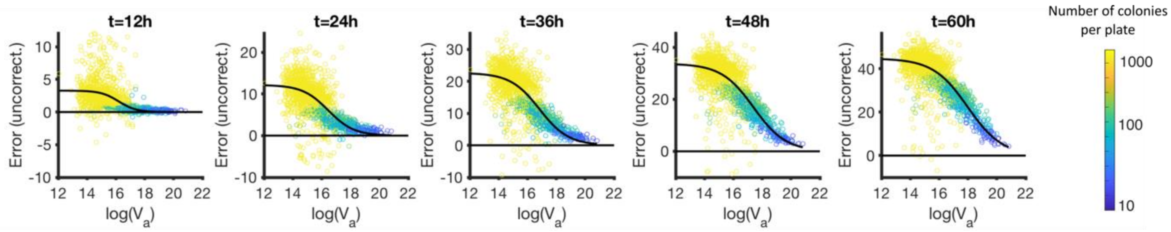

**Supplementary Movie 3:** This movie shows, as in Fig. 8A, the correlation and fitted model of the systematic error obtained after the use of a linear regression with $GR_{max}$ to obtain $t_{app}$. The 0 line thus represents colonies growing at maximal growth rate, and the fit is described in main text. Note the change of the y-axis limits.
